# Tuned inhibitory control of neuronal firing thresholds explains predictive sensorimotor behavior

**DOI:** 10.64898/2026.04.30.721802

**Authors:** Jungryul Ahn, Seolmin Kim, JeongJun Park, Sanghum Woo, Hansem Sohn, Joonyeol Lee

## Abstract

Prior expectations guide sensorimotor behavior when sensory information is uncertain, yet the cellular mechanisms underlying this integration remain elusive. Here, we investigate how priors in sensory motion direction shape neural population dynamics by combining recurrent neural network (RNN) modeling with macaque smooth pursuit behavior and middle temporal area (area MT) electrophysiology. Our RNN model reveals that prior expectations are implemented by elevating firing thresholds in neurons tuned away from the expected direction. This selective inhibition sharpens population tuning and reduces behavioral variability under weak sensory information conditions. We validated this prediction *in vivo*: delta-band local field potentials in area MT, where the phase reflects neural excitability states, exhibited direction-specific phase shifts that scaled with the deviation from the expected direction. These findings demonstrate that prior expectations enhance behavioral reliability through tuned inhibitory control of neuronal excitability, providing a mechanistic link between Bayesian inference and cortical circuit dynamics.

## Introduction

Perception and action are tightly coupled processes that enable organisms to adapt efficiently to dynamically changing environments. Efficient sensorimotor transformation often relies on integrating multiple sources of information—including sensory inputs—to guide motor actions, with each source weighted according to its reliability^1-5^. This process, consistent with *Bayesian inference*, provides a principled computational framework for how predictions derived from prior knowledge are dynamically combined with incoming sensory information. When sensory information is highly reliable, behavior is dominated by current sensory inputs; when it is uncertain or noisy, prior knowledge exerts a stronger influence.

Previous studies have demonstrated neural representations of prior knowledge across multiple brain regions, including the prefrontal cortex, parietal cortex, and even sensory cortices^6-10^. Several studies have further revealed neural mechanisms underlying reliability-based integration, whereby sensory representations are refined by prior expectation. Such expectation-induced enhancement of sensory information can be explained by changes in neural tuning functions for specific stimulus features. In particular, shape changes in population tuning functions—such as sharpening of orientation or motion-direction tuning—have been proposed as key neural substrates for prior-induced perceptual biases or increases in perceptual reliability^5,11-13^. Sharpening of population tuning functions naturally explains enhanced sensory representations and thus provides a population-level mechanism for how prior expectations improve sensory processing. However, the *cellular-level mechanisms* by which prior expectations modulate population tuning functions remain poorly understood.

At the cellular level, sharpening of population tuning may arise from several mechanisms, including tuned modulation of neuronal firing thresholds, regulation of input or response gain, or combinations of these processes. However, directly testing the causal roles of these mechanisms remains experimentally challenging, as current techniques do not allow selective and targeted control of firing thresholds or gains across specific subsets of neurons within a population. Computational modeling— particularly recurrent neural network (RNN) modeling—offers a powerful complementary approach to overcome these limitations^14-16^. RNNs have become increasingly important in neuroscience because they provide full access to network dynamics, connectivity, and unit-level activity, enabling causal tests of mechanistic hypotheses that are difficult or impossible to perform experimentally. Leveraging these advantages, we used RNNs not merely as task-performing models but as controlled testbeds for hypothesis-driven investigation of cellular-level mechanisms underlying prior-dependent modulation of population tuning. Specifically, we focused on the possibility that prior expectations modulate the input–output functions of individual neurons within a population—a form of population-specific gain or threshold control that is currently difficult to isolate experimentally but is accessible through computational models such as RNNs.

In this study, we investigated these cellular-level mechanisms by constraining RNN models with behavioral and neural data recorded from monkeys. Building on prior work that demonstrated expectation-dependent changes in behavior and population tuning—such as tuning sharpening under low sensory reliability^13^, we leveraged these experimental observation to both constrain and evaluate our computational models. We focused on monkeys performing a smooth pursuit eye movement task— a well-established paradigm for probing real-time sensorimotor transformation. By independently manipulating sensory and prior information, we examined how prior knowledge shapes pursuit behavior across different levels of sensory reliability. Using these behavioral data together with neural population direction-tuning functions measured in the middle temporal area (area MT), we found that the RNN model identifies tuned inhibitory control of neuronal firing thresholds as a plausible mechanism for prior-induced behavioral and neural modulation. We further examined this mechanism in a spiking neural network model to assess its biophysical plausibility. We then validated this mechanism by analyzing local field potentials (LFPs) recorded from area MT. These recordings allowed us to test whether sensory cortical activity exhibits signatures of prior-induced inhibitory modulation consistent with the model’s predictions. Together, this integrative approach links behavioral signatures of reliability-weighted Bayesian inference to plausible cellular-level mechanisms in sensory cortex.

## Results

### Prior expectations sharpen sensory cortical population responses and improve pursuit reliability under sensory uncertainty

We trained two monkeys to perform a smooth pursuit eye movement task. Each trial began with fixation on a central dot, followed by the appearance of a moving dot patch. The monkeys were required to track the moving target with their eyes (Fig. 1A). We manipulated two factors: the strength of sensory information and the strength of prior expectation. Sensory information was controlled by stimulus contrast, with high contrast (100%) providing strong sensory evidence and low contrast (10%) providing weak sensory evidence (Fig. 1B). Prior expectation was manipulated through block structure: on a given day, we selected a motion direction as the ‘prior’ direction. In the *no prior* block, motion directions were spaced 120° apart around the prior direction, resulting in high directional uncertainty, whereas in the *prior* block, directions were clustered within 15° of the prior direction, allowing monkeys to develop a strong directional expectation (Fig. 1C). For example, both blocks included the common prior motion direction (e.g., 60°), but its contextual meaning differed: in the *no prior* block, this direction was one of three widely separated possibilities, while in the *prior* block, it was strongly expected. This design yielded four experimental conditions (2 sensory × 2 prior).

**Fig 1:**
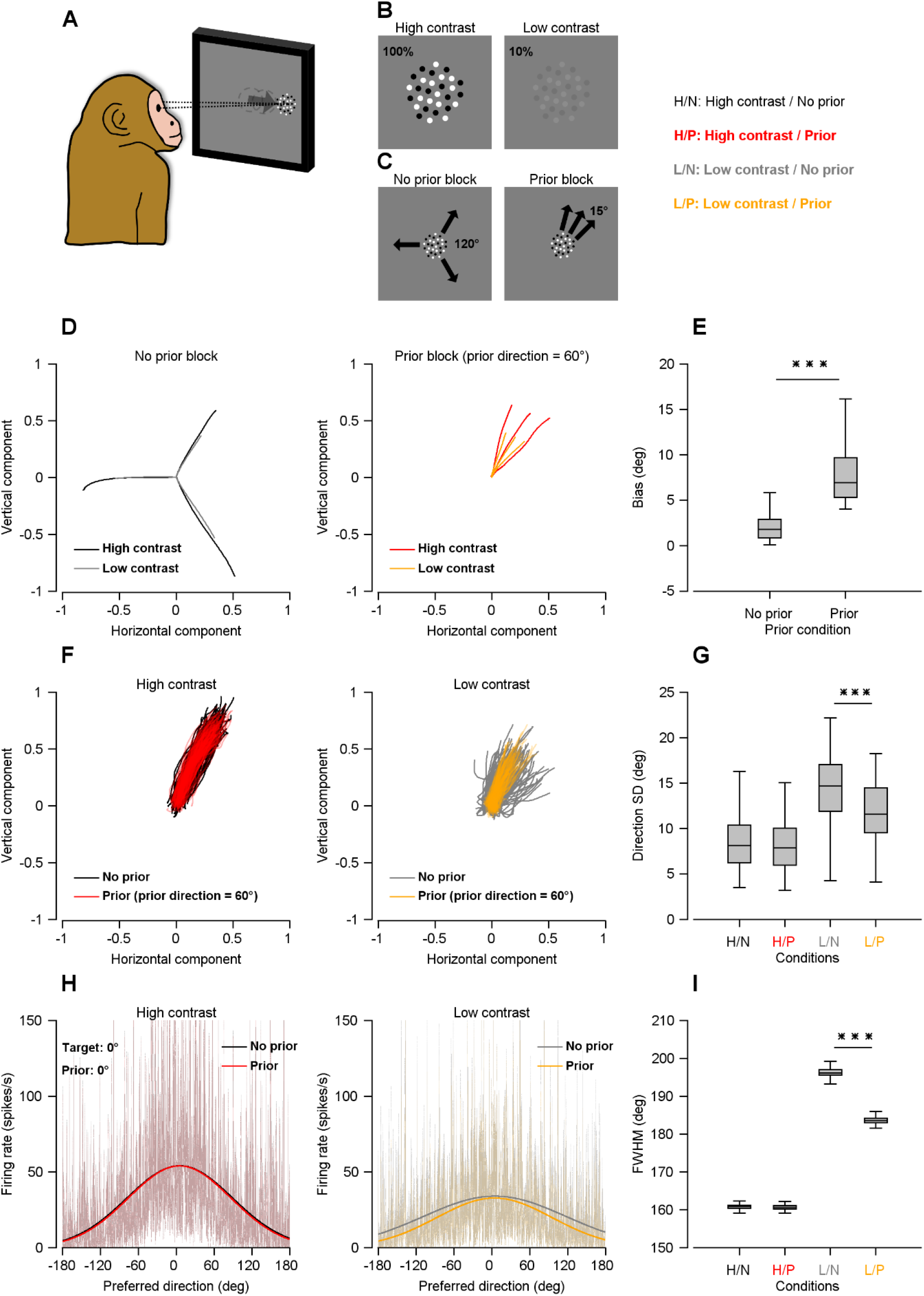
Smooth pursuit eye movement task and behavioral and neural effects of prior expectation. **A** Schematic of the smooth pursuit eye movement task. **B** Target stimulus contrast. Two contrast levels were used: high contrast (100%) and low contrast (8% or 12%, average value: 10%). **C** Manipulation of prior expectation for a specific, example motion direction. The experiment consisted of *prior* and *no prior* blocks. In the *no prior* block, the target moved in one of three directions spaced 120° apart (60°, 180°, and 300°) across trials. In the *prior* block, the target moved in one of three directions spaced 15° apart (60°, 45°, and 75°), sharing a common direction (60°) with the *no prior* block. In the *prior* block, 60° served as the prior direction. **D** Pursuit bias. In the *prior* block, eye movements showed a bias toward the prior direction (60°) when tracking targets moving in nearby directions (45° and 75°). Shown are average pursuit eye trajectories in the *no prior* (left) and *prior* (right) blocks. Horizontal and vertical velocities were normalized by pursuit speed. Black and red traces indicate the eye trajectories for high-contrast stimuli, whereas gray and yellow traces indicate those for low-contrast stimuli. **E** Quantification of pursuit bias. The magnitude of bias toward the prior direction was compared between high and low contrast at each prior condition. In the box plots, the central line indicates the median, the box edges indicate the 25th and 75th percentiles, and the whiskers represent the data range. Paired *t*-test revealed a significantly increased bias under the low-contrast *prior* condition. \*\*\**P* < 0.001. **F** Trial-to-trial variability of pursuit. Each panel shows the monkey’s eye trajectories across 100 trials for the 60° target direction. For both high and low contrast conditions, black and gray traces represent trajectories from the *no prior* block, whereas red and yellow traces represent trajectories from the *prior* block. **G** Quantification of directional variability. The standard deviation (SD) of pursuit direction was computed for each condition. Box plot conventions are as described in (E). Paired *t*-tests for each contrast confirmed that directional variability decreased in the *prior*/low-contrast condition. **H** Reconstructed population direction tuning curves, responding to a motion target with 0° direction. (Left) High-contrast stimulus; (Right) Low-contrast stimulus. Black and gray lines indicate *no prior* conditions and red and yellow lines indicate *prior* conditions. The prior direction was 0°. **I** Quantification of population tuning sharpness. The full width at half maximum (FWHM) of the population tuning curve was measured for each condition. Box plot conventions are as described in (E). Statistical comparisons between *prior* and *no prior* conditions were made using paired *t*-tests for each contrast. \*\*\**P* < 0.001. H/N: high contrast, *no prior*; H/P: high contrast, *prior*; L/N: low contrast, *no prior*; L/P: low contrast, *prior*.

From a Bayesian perspective, combining sensory evidence with prior expectation leads to two key behavioral predictions. First, when sensory information is unreliable, the estimated motion direction should be biased toward the prior, whereas this bias should be minimal when sensory evidence is strong. Second, prior expectations should reduce uncertainty in the estimated direction, resulting in lower trial-to-trial variability, especially under weak sensory conditions. Thus, Bayesian integration predicts both a systematic directional bias and a reduction in behavioral variability when priors are available, with both effects being strongest under low sensory reliability.

Consistent with these predictions, we observed a clear interaction between sensory reliability and prior expectation. In the *no prior* block, both high- and low-contrast stimuli induced similar eye trajectories across the three directions separated by 120° (Fig. 1D). However, in the *prior* block, low-contrast stimuli produced biased trajectories toward the expected 60° direction, especially for nearby stimulus directions of 45° and 75°. Statistically, prior expectation induced a stronger directional bias toward the prior direction under low contrast compared with high contrast (paired *t*-test, *t*(125) = 21.599, *P* = 5.154×10^-44^; Fig. 1E, Supplementary Figure 1A). This indicates that when sensory evidence was weak, prior expectations guided pursuit behavior. In addition, prior expectations reduced trial-to-trial variability in pursuit. This effect was evident under low-contrast conditions, where variability was high without priors but substantially reduced with priors across trials (ANOVA, *F*(3, 500) = 103.634, *P* = 3.513×10^-52^, paired *t*-test, *t*(125) = 14.086, *P* = 1.427×10^-27^; Fig. 1F, G, Supplementary Figure 1B). On the other hand, under high-contrast conditions, pursuit variability was comparable between *prior* and *no prior* blocks (paired *t*-test, *t*(125) = 1.372, *P* = 0.173), suggesting that reliable sensory evidence can override the effect of prior expectation, whereas under weak sensory evidence, priors compensate for reduced precision in tracking behavior.

To investigate the neural correlates of these behavioral effects, we next examined population neural responses in the middle temporal area (area MT) of monkeys, a key region for motion processing. Previous studies have shown that behavioral improvements associated with prior expectations are accompanied by sharpening of neural population tuning, which enhances the precision of sensory representations^11,13^. Consistent with our prior report, and using the same MT recordings, we confirmed this sharpening by reconstructing population neural responses from actual neural recordings (see Methods for detail)^13^. Specifically, prior expectations sharpened population direction tuning in area MT by suppressing responses of neurons whose preferred directions were inconsistent with the expected motion. Under high-contrast conditions, population tuning was similar between *prior* and *no prior* conditions (paired *t*-test, *t*(99) = 0.238, *P* = 0.812; Fig. 1H, I), whereas under low-contrast conditions, population tuning was significantly sharper in the *prior* condition (ANOVA, *F*(3, 396) = 15910.126, *P* < 0.001; paired *t*-test, *t*(99) = 158.728, *P* = 5.481×10^-121^). Thus, neural tuning sharpening was most pronounced when sensory input was noisy (e.g., under low luminance contrast), paralleling the behavioral effects observed under low sensory reliability. We investigated cellular mechanisms for these population-level effects using mechanistic RNN and LFP analyses.

Together, these findings demonstrate that prior expectations influence smooth pursuit eye movements by both biasing estimated motion direction and reducing behavioral variability when sensory inputs are ambiguous. At the neural level, these effects are supported by prior-dependent sharpening of population neural tuning in area MT, consistent with Bayesian integration of sensory evidence and prior knowledge^3,17,18^.

**Supplementary Figure 1:**
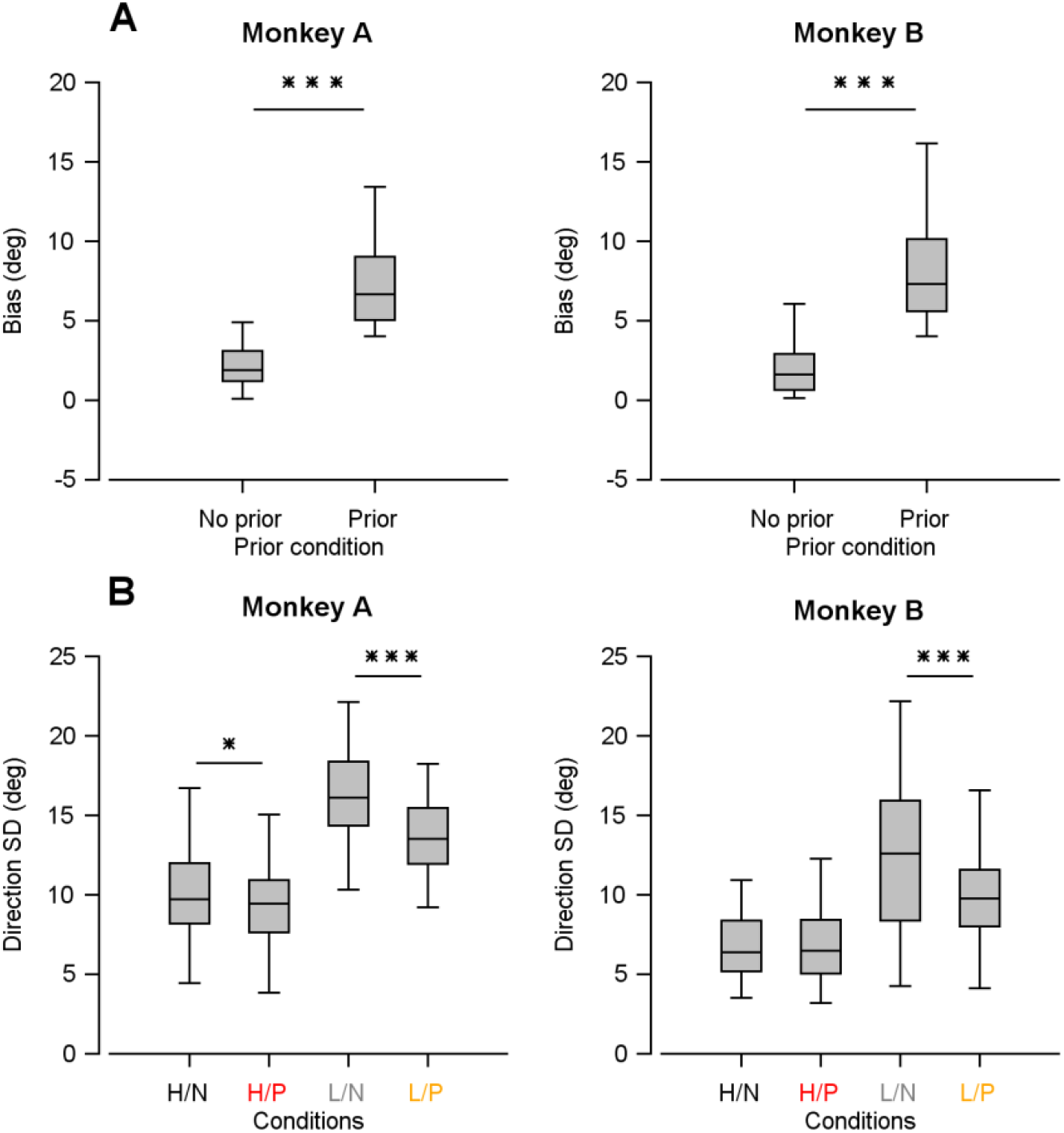
Pursuit bias and variability for each monkey. **A** Quantification of pursuit bias. The magnitude of bias toward the prior direction was measured under each condition. The left panel shows results from monkey A, and the right panel shows results from monkey B. Box plots indicate the median (center line), 25th and 75th percentiles (box edges), and data range (whiskers). Paired *t*-test revealed a significantly increased bias under the low-contrast *prior* condition. \*\*\**P* < 0.001. **B** Quantification of pursuit variability. Directional variability was quantified as the standard deviation (SD) of pursuit direction. The left panel shows results from monkey A, and the right panel shows results from monkey B. Box plot conventions are as described in (A). In the high-contrast condition, monkey A showed a modest but significant decrease in directional variability (\**P* = 0.042), whereas monkey B showed no significant change. In the low-contrast condition, both monkey A and monkey B showed a significant decrease in directional variability. \*\*\**P* < 0.001. H/N: high contrast, *no prior*; H/P: high contrast, *prior*; L/N: low contrast, *no prior*; L/P: low contrast, *prior*.

### Recurrent neural network modeling

#### Emergence of area MT-like direction selectivity in a neural network trained on the smooth pursuit eye movement task

To investigate the potential cellular mechanisms underlying the behavioral effects of prior expectation, we first constructed an RNN model that was designed to reproduce smooth pursuit eye movements (Fig. 2A). The RNN was trained using the target motion of a moving dot patch as the input and the monkey’s actual eye movements as the output. Eye movement traces were obtained from two monkeys across multiple recording days. For training RNN, we used daily averaged eye traces exclusively from the *no prior* condition. This training strategy was adopted to first establish a baseline network capable of generating accurate pursuit behavior and area MT-like direction selectivity in RNN units, independent of prior expectations. This approach allowed us to first isolate the core sensorimotor transformation required for pursuit and then examine how prior-related mechanisms could emerge or be implemented on top of an already functional pursuit network. During training, the model’s performance improved, and the difference between the predicted and actual pursuit traces decreased steadily (Fig. 2B). Following training, the RNN successfully generated pursuit trajectories that closely matched the monkey’s eye movements in all directions of motion (Fig. 2C, D). The individual RNN units comprising the neural network showed robust responses to a specific motion direction (Fig. 2E), and exhibited tuning curves that closely resembled those observed in area MT (Fig. 2F, G). When the neural population activity was aligned with the actual motion direction at 0°, the neural network produced a typical population direction tuning curve (Fig. 2H), similar to the previous study that mimicked the motor cortical responses to arm reaching behavior^19^. This confirmed that the model not only captured pursuit dynamics but also developed neural representations that mirrored empirical observations at both the single-cell and population levels.

**Fig 2:**
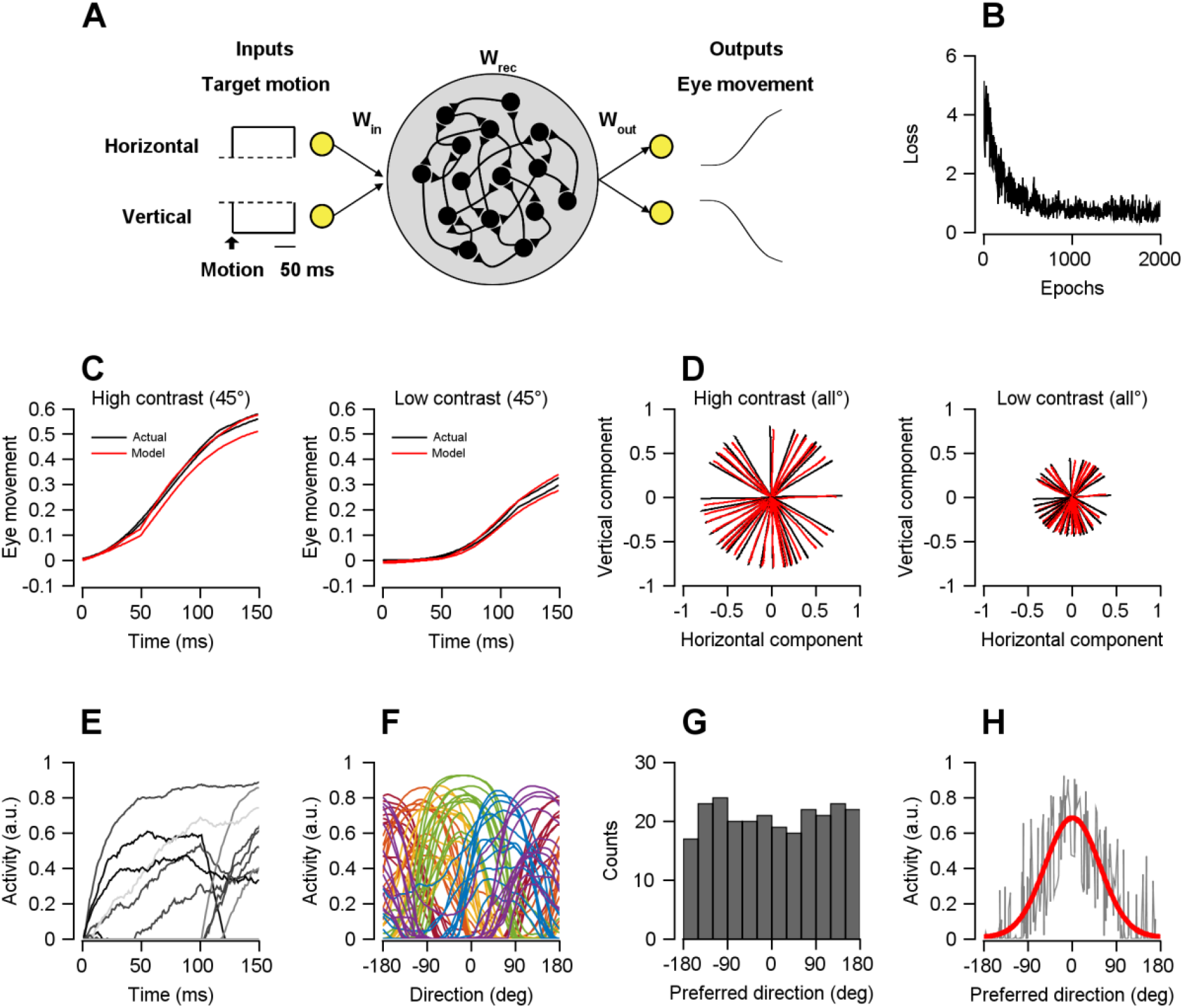
Development of a recurrent neural network (RNN) model for smooth pursuit eye movements. **A** Schematic of the RNN performing the smooth pursuit eye movement task. The network consisted of 250 neurons. The inputs were the horizontal and vertical components of target’s motion direction vector, and the outputs corresponded to the monkey’s normalized eye movements from actual behavioral recordings. During training, the input (*W*_*in*_), recurrent connectivity (*W*_*rec*_), and readout (*W*_*out*_) weights of the RNN were optimized to reproduce the monkey’s eye movements. **B** Change inthe loss function across training iterations, measured as the error between the RNN output and the monkey’s eye movements. **C** Example of eye movement for a target direction of 45°. Black lines represent the monkey’s normalized horizontal and vertical eye velocities, and red lines represent the model-predicted pursuit velocity traces. (Left) High-contrast stimulus condition. (Right) Low-contrast stimulus condition. In the model, sensory contrast was implemented by scaling the amplitude of the input pulse (high: 1, low: 0.1) while preserving the directional ratio defined by *cos(θ)* and *sin(θ)*. **D** Comparison between the monkey’s eye movements (black) and the RNN-predicted eye movements (red) across all target directions. **E** Representative responses of model neurons to target motion in specific directions. Temporal response profiles reflect heterogeneous response latencies across neurons, consistent with the broad distribution of response latencies reported in macaque area MT^20^. **F** Individual tuning curves of model neurons as a function of stimulus direction with their preferred direction color-coded. **G** Distribution of preferred directions across model neurons. **H** Population direction tuning curve for neurons responding to a target direction, here designated as 0°. The red line represents a Gaussian fit to the population tuning curve.

#### Emergent threshold adjustments reproduce prior-dependent effects on pursuit behavior and population neural responses

After establishing a neural network that reproduced both smooth pursuit behavior and MT-like direction selectivity, we next asked how prior expectations could be incorporated into this circuit. For any candidate mechanism to be plausible, it should account not only for behavioral effects on pursuit eye movements but also for the sharpening of neural population tuning observed under prior expectation (Fig. 1H, I). Such modulation could plausibly arise from changes in neuronal excitability, sensitivity to input, or response dynamic range. Accordingly, we hypothesized that prior-dependent effects could be implemented through modulation of neuronal input-output functions, specifically by adjusting the spiking nonlinearities of individual units. To test this hypothesis, we introduced parametric modulation of each unit’s activation function and optimized these parameters to reproduce the observed behavioral bias. Three candidate modulatory parameters were considered (Supplementary Figure 2): (i) threshold, which controls the input level required for activation (horizontal shift); (ii) input gain, which scales sensitivity (slope); and (iii) response gain, which determines the maximum firing rate (amplitude).

**Supplementary Figure 2.**
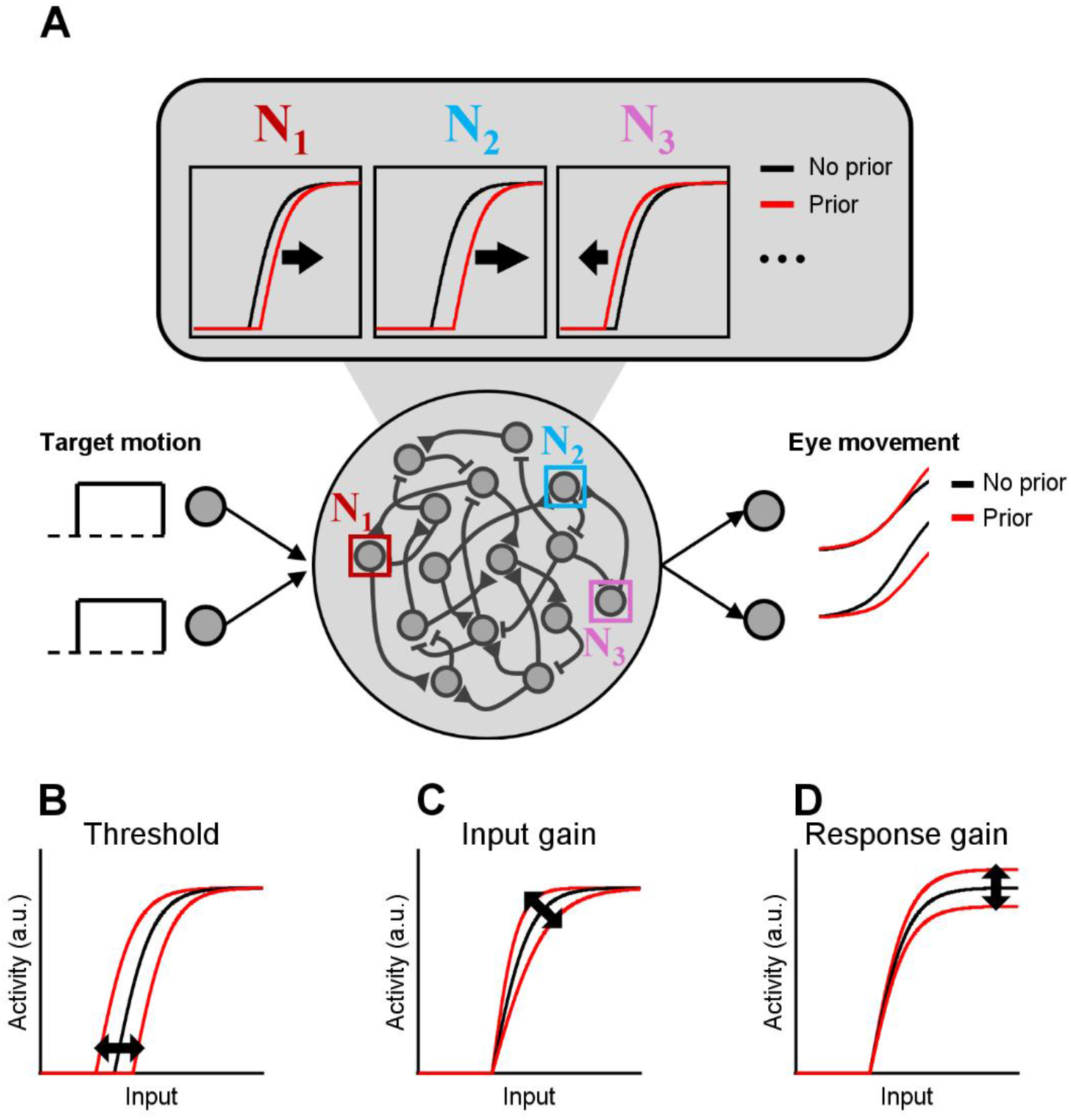
Schematic of parameter modulation in a neuron’s input-output function. **A** The modulatory parameters fitting the eye movement under the prior condition were optimized individually for each neuron, while the input (*W*_*in*_), recurrent connectivity (*W*_*rec*_), and readout (*W*_*out*_) weights were fixed (no training). The black line represents eye traces in the *no prior* condition, and the red line represents eye traces in the *prior* condition. **B** Threshold parameter. Threshold modulation adjusts the minimum input level required for neuronal activation. **C** Input gain parameter. Input gain modulation changes the slope of the input-output function. **D** Response gain parameter. Response gain modulation adjusts the maximum firing rate of the input-output function.

This approach was motivated by prior work showing that modulatory signals in motor cortical circuits can shape complex motor dynamics by training the gain parameters of individual neurons’ input-output functions^21^. More broadly, our approach builds on extensive previous studies showing that neuron-specific threshold or gain modulations can shape population dynamics^22-25^. Crucially, to isolate the effect of prior expectation, synaptic weights were held fixed after the initial training phase (Fig. 2); only the modulatory parameters were subsequently optimized using the monkey’s pursuit bias as the sole objective. As a result, while the reproduction of pursuit bias was a direct outcome of optimization, the reduction in trial-to-trial pursuit variability and the sharpening of neural population tuning were not explicitly constrained. These emergent properties therefore serve as independent model validations, arising only if the identified modulatory mechanism provides a plausible account of the experimental data.

To identify which parameters drove these effects, we optimized each parameter individually while holding the others fixed. Threshold modulation alone was sufficient to reproduce both the behavioral and neural signatures of the experimental observations. At the population level, threshold modulation sharpened direction tuning by selectively suppressing neurons whose preferred directions were distant from the prior (Fig. 3A). Under high-contrast conditions, population tuning curves were similar regardless of whether the threshold modulation was applied (paired *t*-test, *t*(99) = 1.591, *P* = 0.115; Fig. 3B). Under low-contrast conditions, however, threshold modulation produced a pronounced sharpening of the tuning curve (ANOVA, *F*(3, 396) = 9670.249, *P* < 0.001; paired *t*-test, *t*(99) = 62.735, *P* = 1.603×10^-81^). This selective sharpening reproduced the tuning changes observed in area MT, even though preferred directions of individual units were not explicitly constrained during RNN training. A similar pattern emerged for pursuit variability. Trial-to-trial variability did not differ between *prior* and *no prior* conditions under high-contrast (paired *t*-test, *t*(99) = 1.929, *P* = 0.057; Fig. 3D, E), but was significantly reduced by prior expectation under low-contrast (ANOVA, *F*(3, 396) = 6796.226, *P* < 0.001; paired *t*-test, *t*(99) = 58.420, *P* = 1.545 × 10^−78^), consistent with behavioral observations in monkeys (Fig. 1G). In the behavioral data, pursuit variability (mean ± SD) decreased from 14.283 ± 3.988 in the low-contrast, *no prior* condition to 11.649 ± 3.139 in the low-contrast, *prior* condition. The threshold-modulated model reproduced a comparable reduction, yielding values of 16.312 ± 0.956 and 11.034 ± 0.451, respectively. Importantly, the optimized thresholds were not randomly distributed across neurons but exhibited a structured pattern dependent on the prior direction: thresholds increased systematically with the angular distance between the prior direction and each neuron’s preferred direction (Fig. 3C). Neurons tuned away from the prior thus required stronger input to become active, reflecting an emergent, structured prior-dependent modulation of neuronal excitability.

**Fig 3:**
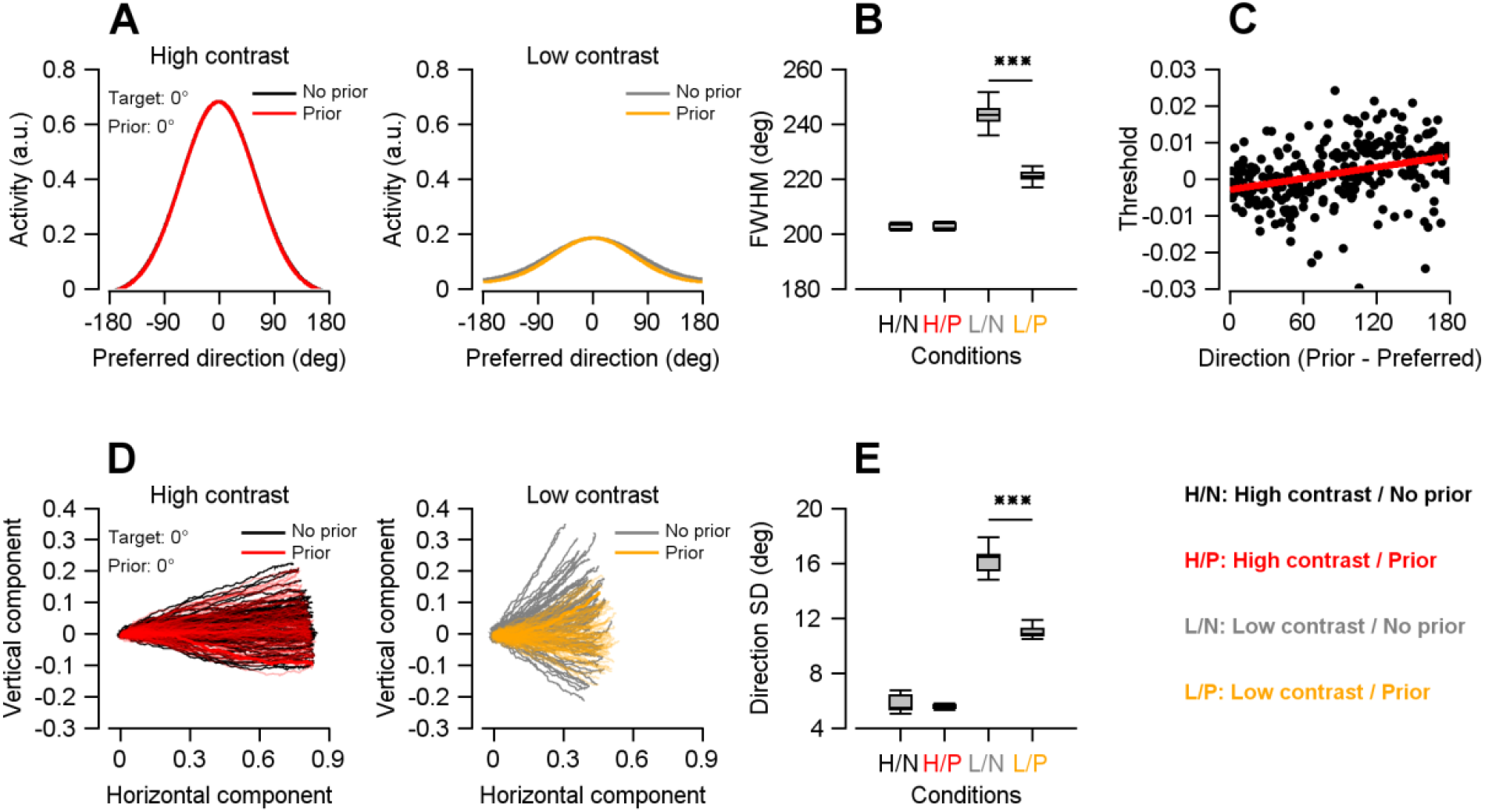
Emergent threshold adjustments. **A** Population direction tuning curves. Population responses are shown for a target direction of 0°. (Left) High contrast stimulus; (Right) Low contrast stimulus. Black and gray lines indicate *no prior* conditions and red and yellow lines indicate *prior* conditions. The prior direction was 0°. **B** Quantification of population tuning sharpness. The full width at half maximum (FWHM) of the population tuning curve was measured for each condition. In the box plots, the central line indicates the median, the box edges indicate the 25th and 75th percentiles, and the whiskers represent the data range. Statistical comparisons between *prior* and *no prior* conditions were made using paired *t*-tests for each contrast. \*\*\**P* < 0.001. **C** Changes in the threshold parameters of individual neurons. The scatter plot shows the relationship between the angular difference between the prior direction and each neuron’s preferred direction and the corresponding threshold value. Each point represents a single neuron, and the red line indicates the linear regression fit. **D** Trial-to-trial variability in pursuit. Eye trajectories from 100 trials are shown for the 0° target direction. For both contrast conditions, high(left) and low(right), black and gray traces represent trajectories from the *no prior* block, whereas red and yellow traces represent trajectories from the *prior* block. The prior direction was 0°. **E** Quantification of directional variability. The standard deviation (SD) of pursuit direction was computed for each condition. Box plot conventions are as described in (B). Paired *t*-tests for each contrast confirmed that directional variability decreased under the low-contrast *prior* condition. \*\*\**P* < 0.001. H/N: high contrast, *no prior*; H/P: high contrast, *prior*; L/N: low contrast, *no prior*; L/P: low contrast, *prior*.

However, input gain and response gain modulation failed to reproduce either the behavioral or neural features observed experimentally. In both cases, the optimized gain parameters decreased monotonically as the angular distance from the prior direction increased (Supplementary figure 3C, 4C). Under both high- and low-contrast conditions, population tuning curves did not differ significantly between the *prior* and *no prior* conditions (input gain modulation: high contrast, paired *t*-test, *t*(99) = 1.377, *P* = 0.172; low contrast, *t*(99) = 1.727, *P* = 0.087; response gain modulation: high contrast, paired *t*-test, *t*(99) = 1.785, *P* = 0.077; low contrast, *t*(99) = 1.931, *P* = 0.056; Supplementary figure 3B, 4B). Additionally, neither input gain nor response gain modulation produced a significant reduction in trial-to-trial pursuit variability under low-contrast conditions (input gain modulation: high contrast, paired *t*-test, *t*(99) = 1.932, *P* = 0.056; low contrast, *t*(99) = 1.838, *P* = 0.069; response gain modulation: high contrast, paired *t*-test, *t*(99) = 1.418, *P* = 0.159; low contrast, *t*(99) = 0.920, *P* = 0.360; Supplementary figure 3E, 4E).

With respect to pursuit bias, prior expectation induced a stronger directional bias toward the prior direction under low-contrast than high-contrast conditions across all modulation parameters (threshold, input gain, and response gain modulation; threshold modulation: paired *t*-test, *t*(99) = 30.601, *P* = 2.888×10^-52^; input gain modulation: paired *t*-test, *t*(99) = 11.462, *P* = 7.325×10^-20^; response gain modulation: paired *t*-test, *t*(99) = 5.372, *P* = 5.179× 10^-7^; Supplementary figure 5). This result is expected, because pursuit bias was explicitly included in the training objective when optimizing the input-output function parameters using monkey behavioral data.

Again, neither the reduction in trial-to-trial pursuit variability nor the sharpening of neural population tuning was included in the training objective. The modulatory parameters were optimized solely to reproduce biased mean pursuit trajectories for individual motion directions. Nevertheless, only threshold modulation gave rise to both reduced pursuit variability under low-contrast *prior* conditions and sharpening of population tuning curve, consistent with the behavioral and neural experiment observations (Fig. 1). In sum, these results indicate that threshold modulation, rather than input or response gain modulation, serves as the primary cellular mechanism underlying the improvement in behavioral reliability induced by prior expectation.

**Supplementary Figure 3:**
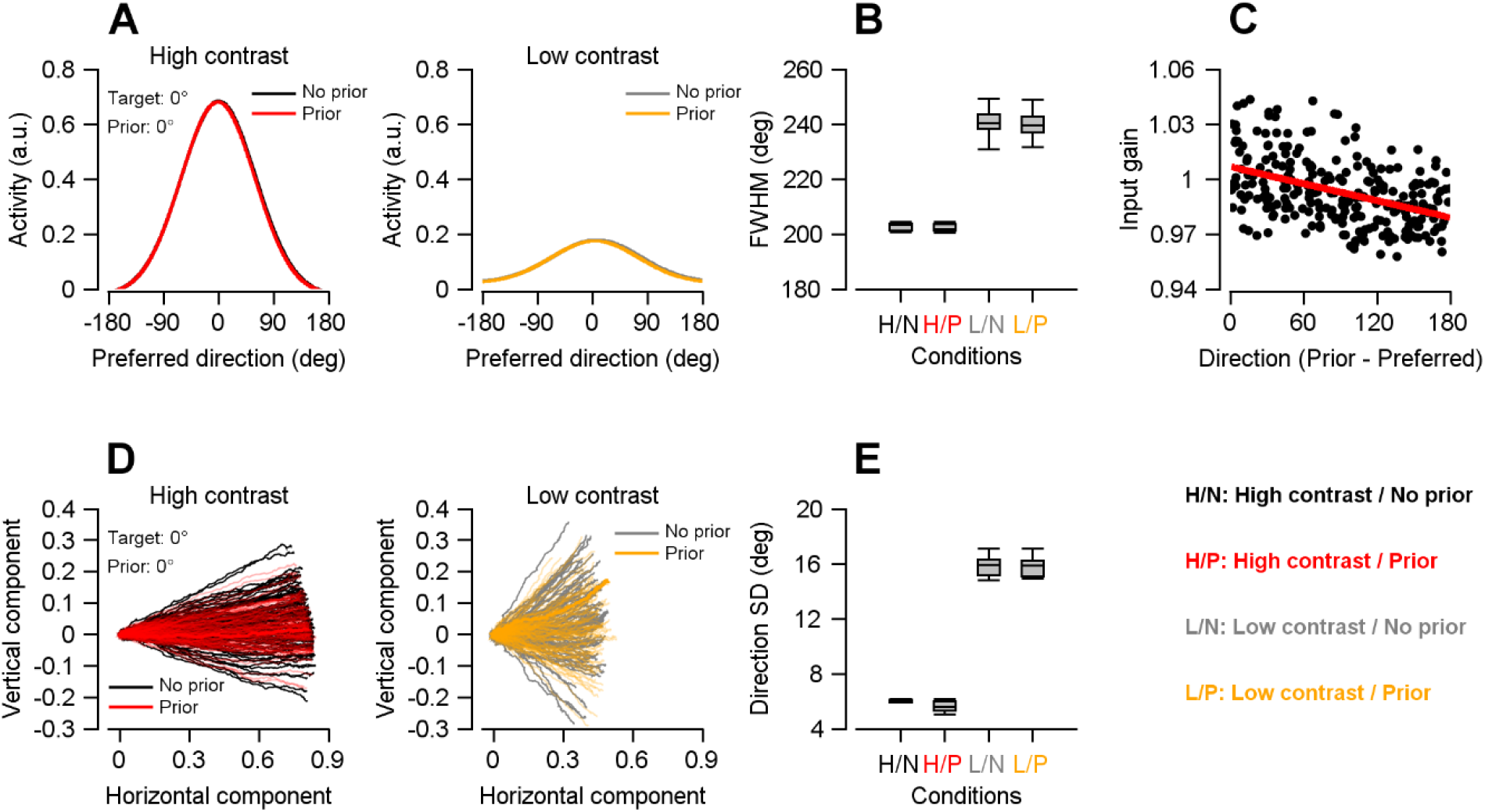
Emergent input gain adjustments. **A** Population direction tuning curves. Population responses are shown for a target direction of 0°. (Left) High contrast stimulus; (Right) Low contrast stimulus. Black and gray lines indicate *no prior* conditions and red and yellow lines indicate *prior* conditions. The prior direction was 0°. **B** Quantification of population tuning sharpness. The full width at half maximum (FWHM) of the population tuning curve was measured for each condition. In the box plots, the central line indicates the median, the box edges indicate the 25th and 75th percentiles, and the whiskers represent the data range. **C** Changes in the input gain parameters of individual neurons. The scatter plot shows the relationship between the angular difference between the prior direction and each neuron’s preferred direction and the corresponding input gain value. Each point represents a single neuron, and the red line indicates the linear regression fit. **D** Trial-to-trial variability in pursuit. Eye trajectories from 100 trials are shown for the 0° target direction. For both contrast conditions, high(left) and low(right), black and gray traces represent trajectories from the *no prior* block, whereas red and yellow traces represent trajectories from the *prior* block. The prior direction was 0°. **E** Quantification of directional variability. The standard deviation (SD) of pursuit direction was computed for each condition. Box plot conventions are as described in (B). H/N: high contrast, *no prior*; H/P: high contrast, *prior*; L/N: low contrast, *no prior*; L/P: low contrast, *prior*.

**Supplementary Figure 4:**
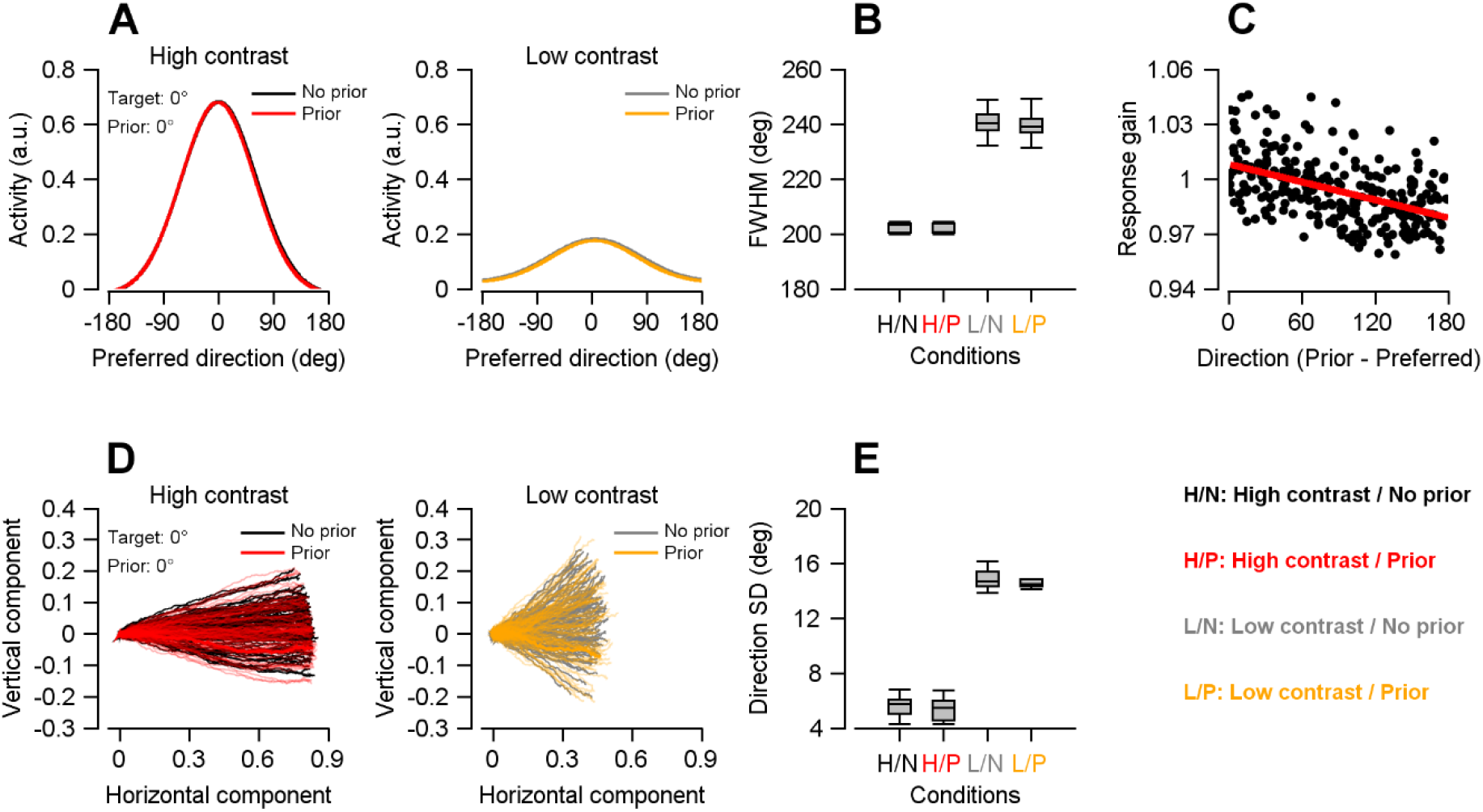
Emergent response gain adjustments. **A** Population direction tuning curves. Population responses are shown for a target direction of 0°. (Left) High contrast stimulus; (Right) Low contrast stimulus. Black and gray lines indicate *no prior* conditions and red and yellow lines indicate *prior* conditions. The prior direction was 0°. **B** Quantification of population tuning sharpness. The full width at half maximum (FWHM) of the population tuning curve was measured for each condition. In the box plots, the central line indicates the median, the box edges indicate the 25th and 75th percentiles, and the whiskers represent the data range. **C** Changes in the response gain parameters of individual neurons. The scatter plot shows the relationship between the angular difference between the prior direction and each neuron’s preferred direction and the corresponding response gain value. Each point represents a single neuron, and the red line indicates the linear regression fit. **D** Trial-to-trial variability in pursuit. Eye trajectories from 100 trials are shown for the 0° target direction. For both contrast conditions, high(left) and low(right), black and gray traces represent trajectories from the *no prior* block, whereas red and yellow traces represent trajectories from the *prior* block. The prior direction was 0°. **E** Quantification of directional variability. The standard deviation (SD) of pursuit direction was computed for each condition. Box plot conventions are as described in (B). H/N: high contrast, *no prior*; H/P: high contrast, *prior*; L/N: low contrast, *no prior*; L/P: low contrast, *prior*.

**Supplementary Figure 5:**
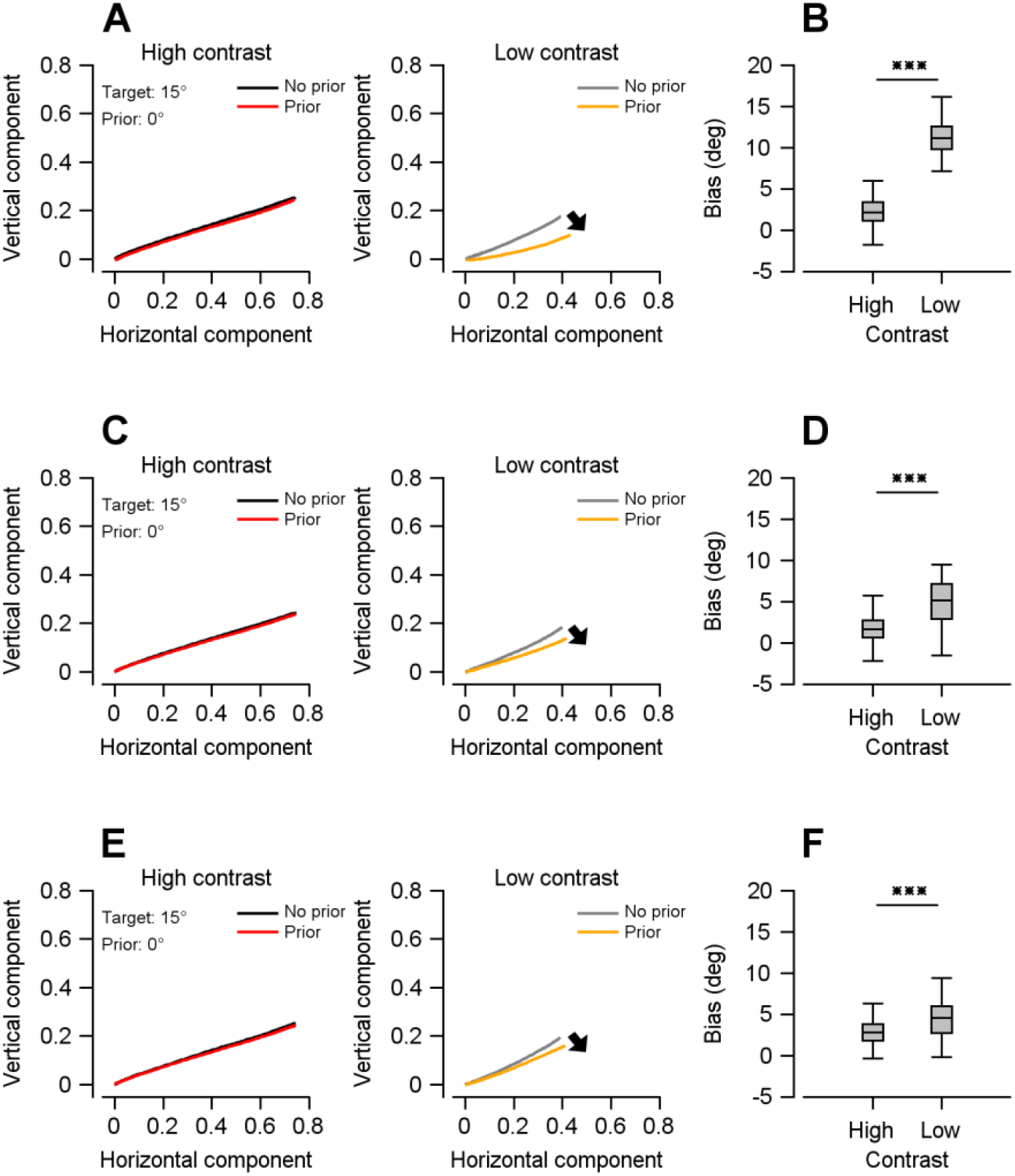
Pursuit bias across different input–output function modulations. **A** Threshold modulation. The model’s smooth pursuit eye trajectories are shown for a target direction of 15°. (Left) High contrast stimulus; (Right) Low contrast stimulus. Black and red traces indicate the eye trajectories for high-contrast stimuli, whereas gray and yellow traces indicate those for low-contrast stimuli. The prior direction was 0°. The black arrow highlights that, under the low-contrast *prior* condition, the eye trajectory was biased toward the prior direction (0°). **B** Quantification of pursuit bias. The magnitude of bias toward the prior direction was compared between *prior* and *no prior* conditions at each stimulus contrast. Here, bias was defined as the deviation of pursuit direction toward the prior direction relative to the *no prior* condition. In the box plots, the central line indicates the median, the box edges indicate the 25th and 75th percentiles, and the whiskers represent the data range. Paired *t*-test revealed a significantly increased bias under the low-contrast *prior* condition. \*\*\**P* < 0.001. **C-D** Input gain modulation. **E-F** Response gain modulation.

#### Predicted neural and behavioral changes during spontaneous activity emerging from the threshold modulation

Prior knowledge, acquired through repeated experience, can become embedded within the neural network. As shown in area MT neurons previously^13^ (Supplementary figure 6A), the neural representation of this prior persists even in the absence of external sensory stimuli (the spontaneous activity period). To simulate this, we injected Gaussian noise into the trained RNN during the period corresponding to the fixation window. This noisy input allows the network to operate in a baseline state, revealing how intrinsic prior information generates directional biases.

We found that the threshold modulation of the input-output function showed the neural and behavioral signatures of the prior expectation while other parameter modulations (input gain, response gain) did not. When direction-selective threshold modulation was applied, noise injection revealed that neurons with preferred directions farther from the prior direction (0°) showed a greater reduction in spontaneous activity, resulting in direction-selective tuning centered on the prior direction (Fig. 4A). These neural changes appeared to produce spontaneous eye drifts toward the prior direction even in the absence of sensory input (Fig. 4B). These model predictions were consistent with experimental observations in monkeys: spontaneous activity in the *prior* condition was tuned to the prior direction when compared with the *no prior* condition (Supplementary figure 6A, B).

**Fig 4:**
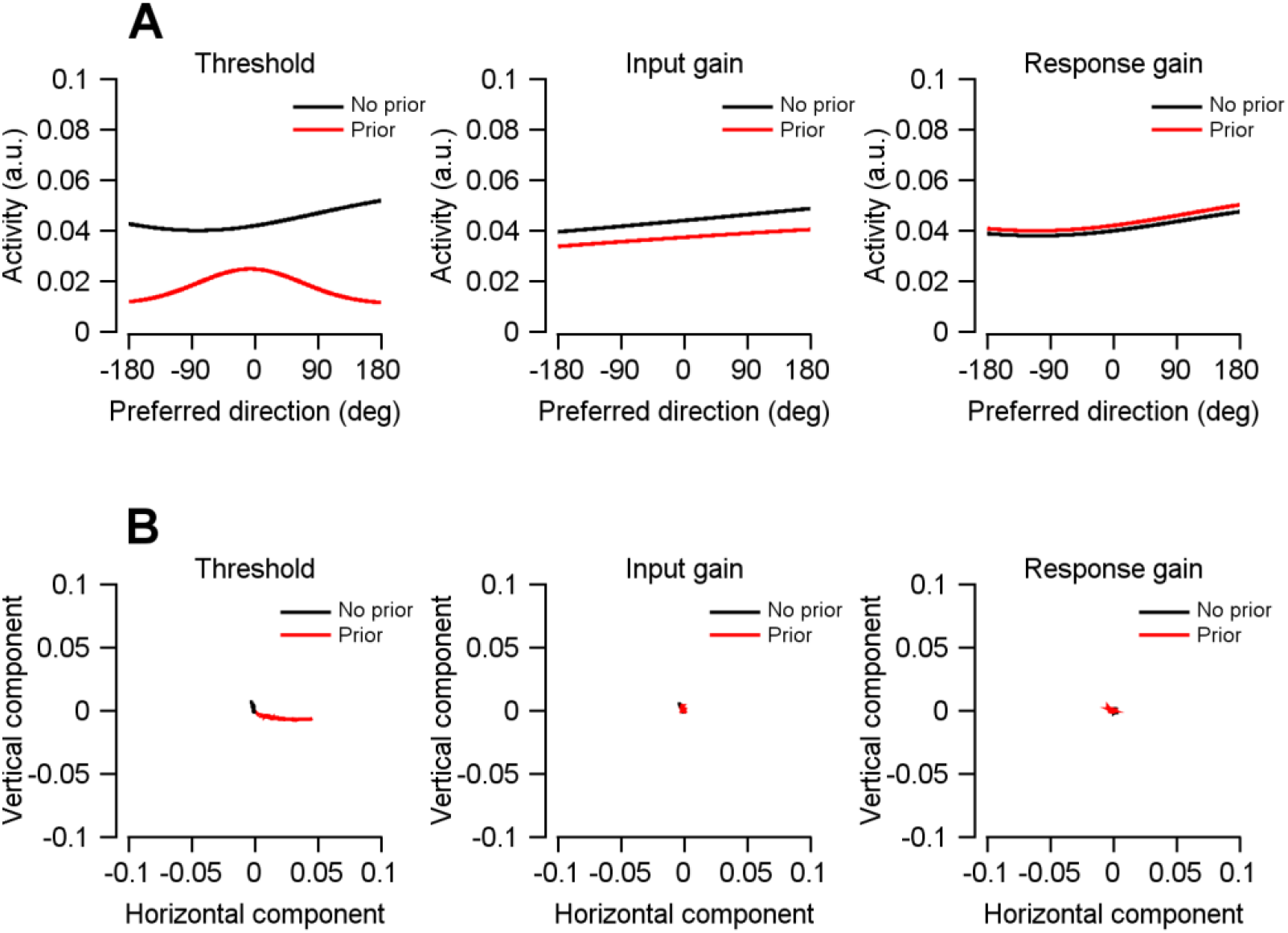
Spontaneous neural population activity and behavior through noisy input injection into the trained neural network. **A** Population directional tuning curves in response to noisy input. (Left) Threshold, (Middle) Input gain, and (Right) Response gain. Black and red lines represent the *no prior* and *prior* conditions, respectively. The prior direction was 0°. **B** Pursuit bias. Model’s smooth tracking eye trajectories for noisy input. Black and red lines represent the *no prior* and *prior* conditions, respectively. The prior direction was 0°.

In contrast, modulation of input gain or response gain failed to reproduce these neural and behavioral effects. Under these conditions, noise injections did not generate direction-selective tuning in spontaneous neural activity, nor did it produce systematic behavioral biases toward the prior direction. These results indicate that input gain and response gain modulation are insufficient to intrinsically encode prior expectations in spontaneous activity.

In summary, our RNN modeling demonstrates that prior expectations can be implemented through increased thresholds in neuronal input-output functions. This mechanism operates by reducing the baseline excitability of neurons misaligned with the prior direction, thereby sharpening neural population activity, suppressing behavioral variability, and biasing pursuit behavior toward the expected direction. Consequently, it enhances behavioral accuracy and reliability under uncertain sensory conditions.

**Supplementary figure 6:**
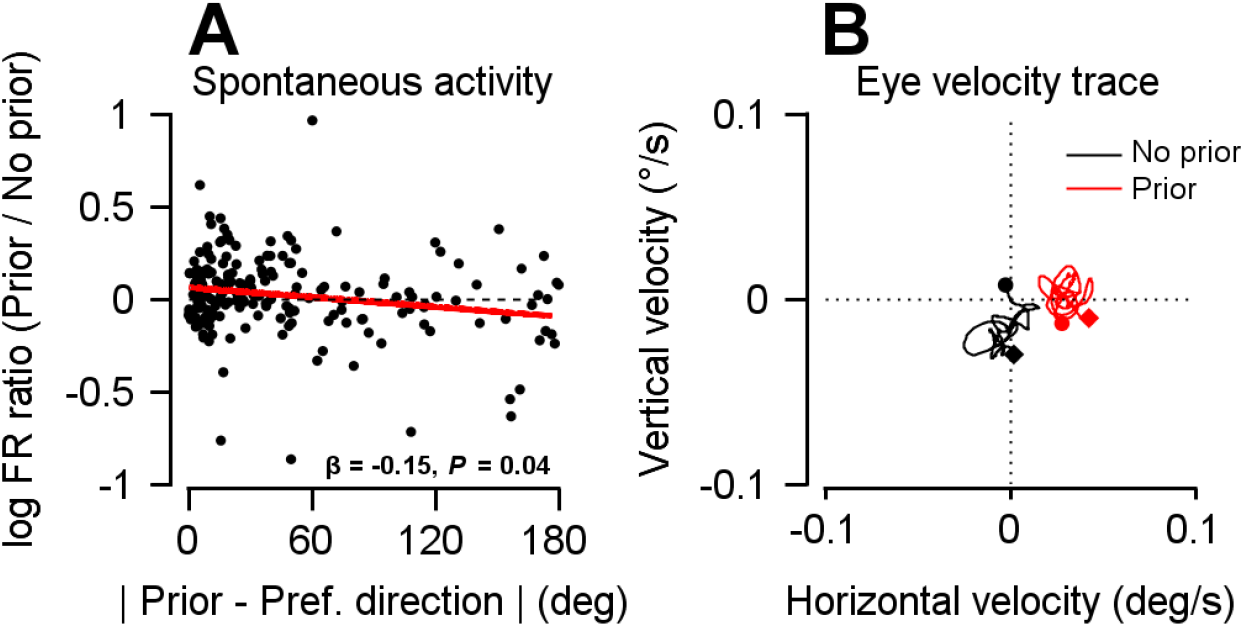
Spontaneous neural population activity and behavior in monkeys under different prior conditions. **A** Firing rate log ratio (*prior vs. no prior*) during the spontaneous activity period (−400 to 0 ms) in area MT neurons recorded from two monkeys, plotted as a function of the angular difference between the prior direction and each neuron’s preferred direction. Each point represents an individual neuron. The red line indicates a linear regression fit (β is regression coefficient, significance expressed as *P* value). **B** Eye velocity traces during the fixation period (−400 to 0 ms from motion onset) under *no prior* and *prior* blocks. Smooth pursuit eye movement traces before motion onset reveal prior-dependent biases in baseline eye velocity. Data from one monkey; eye velocity was rotated so that 0° corresponds to the prior direction. Figures adapted from Supplementary Figure 9 of Park et al. (2023)^13^.

#### Standard weight-based RNN optimization failed to reproduce prior-induced modulation of population direction tuning

Our modeling approach has focused on modulation of neuronal input-output functions, rather than changes in synaptic connectivity, as a mechanism for implementing the effects of prior expectation. To directly compare this approach with more conventional frameworks, we next tested a standard weight-based RNN model in which prior information was provided explicitly as an additional input signal, and its effects were learned through synaptic weight optimization (Fig. 5A). Using the same pursuit bias measurements as in the input-output function parameter optimization, we optimized the input, recurrent, and readout weights to reproduce the monkey’s pursuit bias behavior, while keeping the input-output function parameters fixed across prior conditions. This approach follows standard RNN frameworks in which contextual modulation is implemented through weight-based adjustments rather than changes in neuronal input-output properties^15,26^.

**Fig 5:**
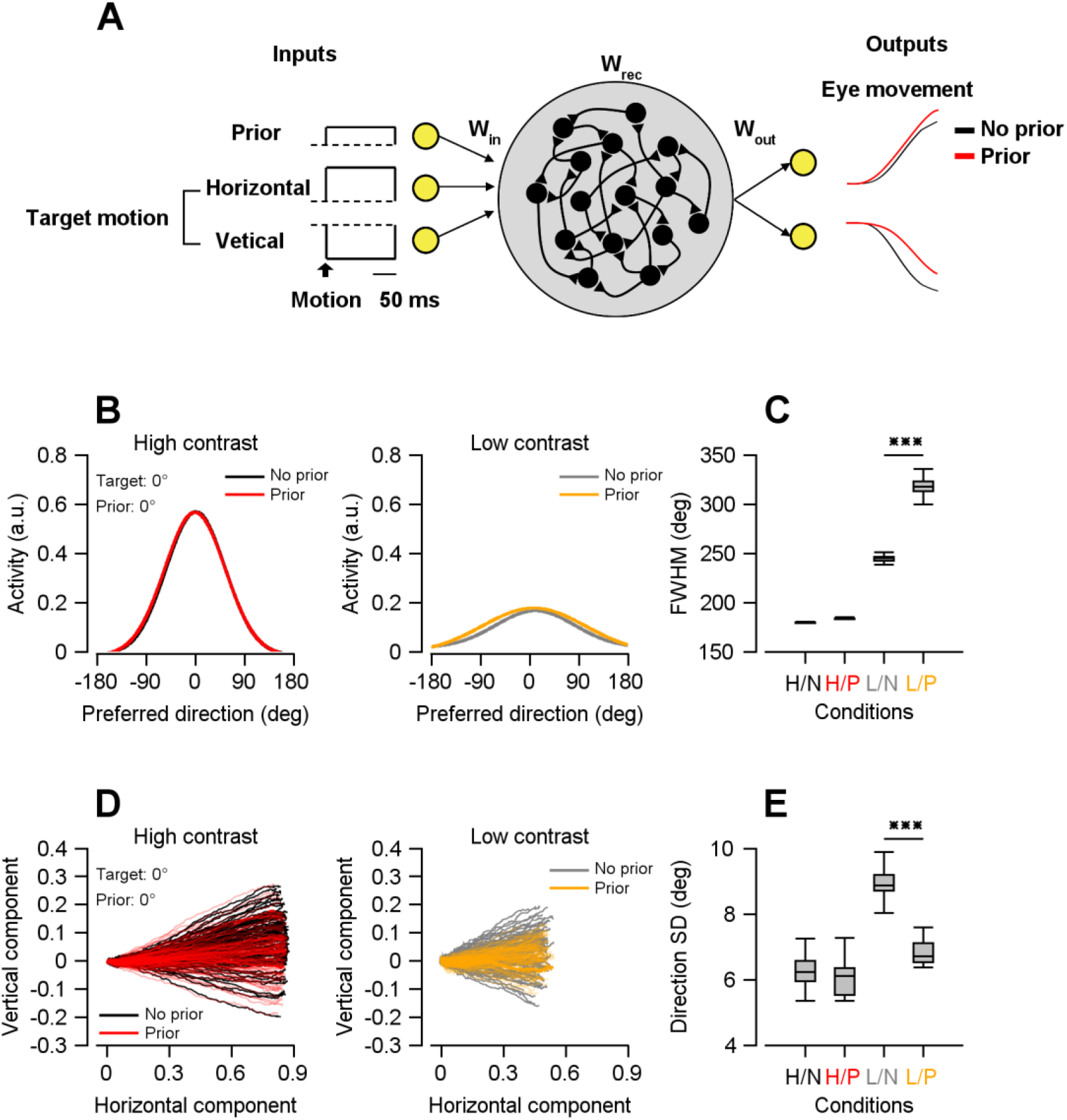
Effects of applying prior input on neural tuning and pursuit behavior in the RNN model. **A** Schematic diagram of an RNN with prior input. Network inputs consisted of a prior signal and the horizontal and vertical components of target’s motion direction vector. The network outputs corresponded to the monkey’s normalized eye movements obtained from behavioral recordings. During training, the input (*W*_*in*_), recurrent connectivity (*W*_*rec*_), and readout (*W*_*out*_) weights of the RNN were optimized to reproduce the monkey’s eye movements. Black traces indicate normalized horizontal and vertical eye velocities in the *no prior* condition, and red traces indicate those in the *prior* condition. **B** Population direction tuning curves. Population responses are shown for a target direction of 0°. (Left) High contrast stimulus; (Right) Low contrast stimulus. Black and gray lines indicate *no prior* conditions and red and yellow lines indicate *prior* conditions. The prior direction was 0°. **C** Quantification of population tuning sharpness. The full width at half maximum (FWHM) of the population tuning curve was measured for each condition. In the box plots, the central line indicates the median, the box edges indicate the 25th and 75th percentiles, and the whiskers represent the data range. Statistical comparisons between *prior* and *no prior* conditions were made using paired *t*-tests for each contrast. \*\*\**P* < 0.001. **D** Trial-to-trial variability in pursuit. Eye trajectories from 100 trials are shown for the 0° target direction. For both contrast conditions, high(left) and low(right), black and gray traces represent trajectories from the *no prior* block, whereas red and yellow traces represent trajectories from the *prior* block. The prior direction was 0°. **E** Quantification of directional variability. The standard deviation (SD) of pursuit direction was computed for each condition. Box plot conventions are as described in (B). Paired *t*-tests for each contrast confirmed that directional variability decreased under the low-contrast *prior* condition. \*\*\**P* < 0.001. H/N: high contrast, *no prior*; H/P: high contrast, *prior*; L/N: low contrast, *no prior*; L/P: low contrast, *prior*.

The weight-optimized model successfully reproduced the behavioral pursuit bias, indicating that synaptic weight changes alone can account for the prior-induced shift in pursuit behavior (paired *t*-test, *t*(99) = 18.363, *P* = 1.195 × 10^−33^; Supplementary figure 7). In addition, the model reproduced a reduction in inter-trial pursuit variability under low-contrast conditions, despite this effect not being explicitly included in the training objective (ANOVA, *F*(3, 396) = 429.045, *P* = 5.198×10^-124^; high contrast: paired *t*-test, *t*(99) = 1.777, *P* = 0.0786; low contrast: *t*(99) = 49.345, *P* = 1.613 × 10^−71^; Fig. 5D, E).

However, the model failed to reproduce a critical neural signature of prior expectation: the sharpening of population tuning functions selectively under low-contrast conditions. This tuning sharpening is consistently observed in the monkey’s neurophysiological data and was not included in the training objective. Instead, introducing the prior input led to a broadening of population tuning curves accompanied by an overall increase in neural responsiveness under low-contrast conditions (ANOVA, *F*(3, 396) = 660.331, *P* = 1.128×10^-153^; high contrast: *t*(99) = 0.667, *P* = 0.506; low contrast: *t*(99) = 90.622, *P* = 4.633 × 10^−97^; Fig. 5B, C). Although this pattern resembles certain attention-related effects, such as response increase, it contrasts with the direction-specific tuning sharpening reported in area MT^27-29^. These findings indicate that weight-based optimization driven solely by a prior input is insufficient to explain how prior expectations shape population activity in sensory cortical circuits.

**Supplementary Figure 7:**
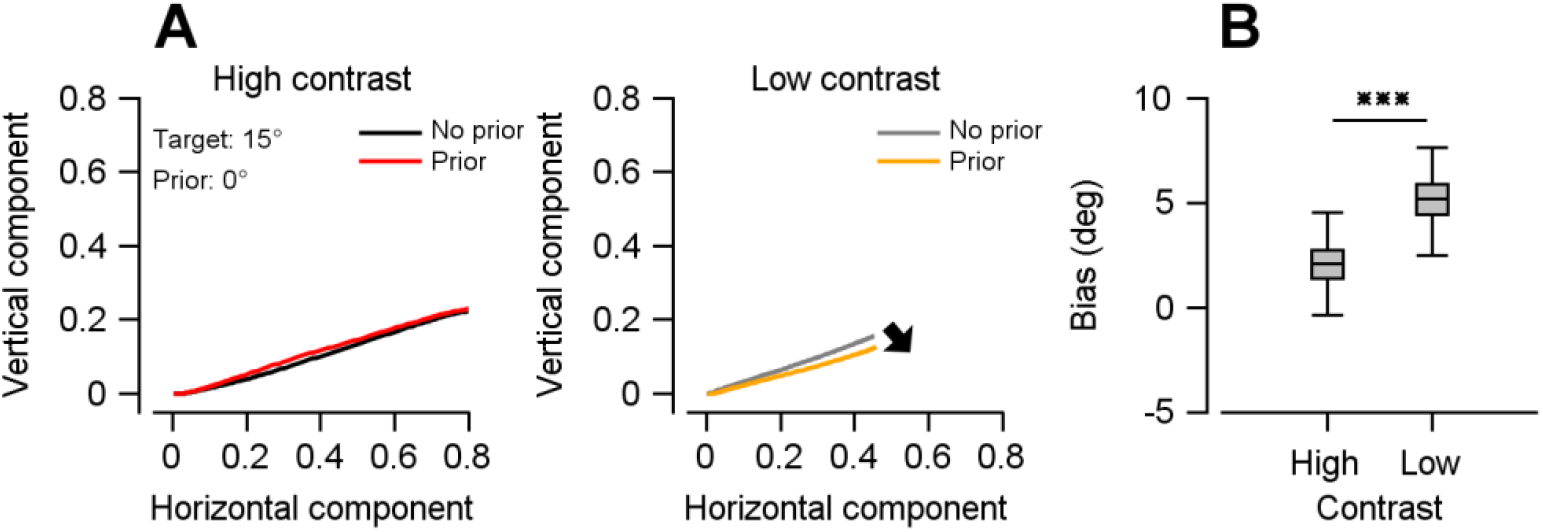
Pursuit bias reproduced by a weight-optimized RNN model. **A** The model’s smooth pursuit eye trajectories are shown for a target direction of 15°. (Left) High contrast stimulus; (Right) Low contrast stimulus. Black and red traces indicate the eye trajectories for high-contrast stimuli, whereas gray and yellow traces indicate those for low-contrast stimuli. The prior direction was 0°. The black arrow highlights that, under the low-contrast *prior* condition, the eye trajectory was biased toward the prior direction (0°). **B** Quantification of pursuit bias. The magnitude of bias toward the prior direction was compared between *prior* and *no prior* conditions at each stimulus contrast. In the box plots, the central line indicates the median, the box edges indicate the 25th and 75th percentiles, and the whiskers represent the data range. Paired *t*-test revealed a significantly increased bias under the low-contrast *prior* condition. \*\*\**P* < 0.001.

#### Tuned inhibitory input to membrane potential accounts for prior-dependent modulation in spiking neural network

The threshold modulation identified in the RNN corresponds, at the biophysical level, to systematic shifts in membrane potential driven by inhibitory input. To directly test this interpretation, we constructed a spiking neural network model in which membrane potential was explicitly modulated by direction-selective inhibitory input. The network consisted of 500 leaky integrate-and-fire (LIF) neurons distributed along a circular axis of preferred motion directions, organized with structured recurrent connectivity following a bump attractor architecture^30,31^. When a directional stimulus was presented, neurons responded with spike trains determined by the match between the stimulus direction and their preferred directions, giving rise to a population activity “bump” centered on the stimulus. The represented motion direction was decoded from spiking responses using population vector decoding, allowing us to directly link neural population dynamics to predicted pursuit behavior (Fig. 6A).

**Fig 6:**
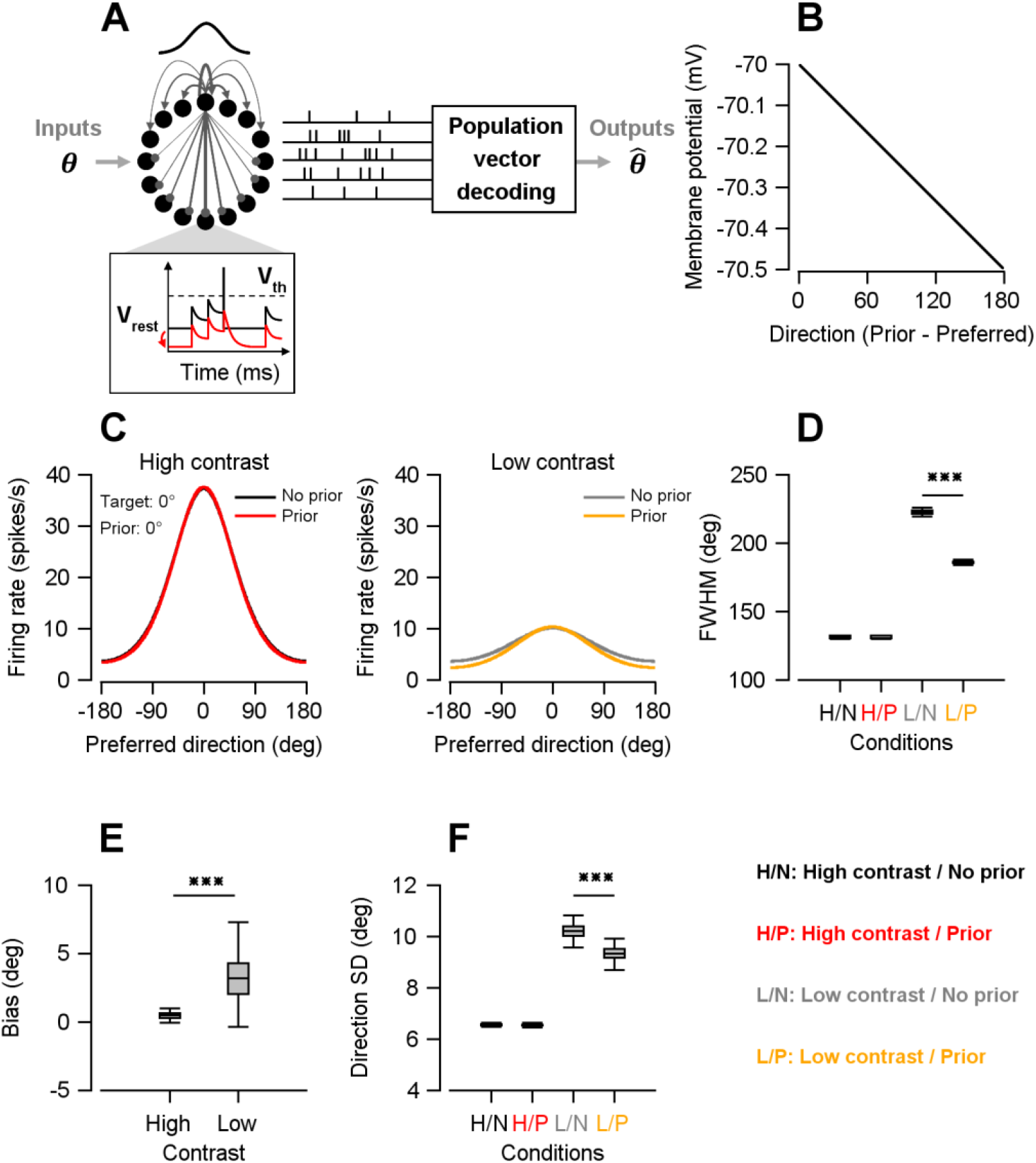
Inhibitory modulation of membrane potential accounts for prior-dependent neural and behavioral changes. **A** Schematic of a spiking neural network model based on leaky integrate-and-fire (LIF) neurons. The network consisted of 500 neurons arranged along a circular axis of preferred motion directions. Directional stimuli evoked population activity forming a localized “bump” centered on the stimulus direction. The represented motion direction was decoded from population activity using population vector decoding. **B** Direction-selective inhibitory modulation. A direction-dependent decrease in membrane potential was applied such that neurons whose preferred directions were farther from the prior direction received stronger inhibitory input. This modulation reduced neuronal excitability in a structured manner, effectively implementing prior-dependent changes in network activity. **C** Population direction tuning curves. Population responses are shown for a target direction of 0°. (Left) High contrast stimulus; (Right) Low contrast stimulus. Black and gray lines indicate *no prior* conditions and red and yellow lines indicate *prior* conditions. The prior direction was 0°. **D** Quantification of population tuning sharpness. The full width at half maximum (FWHM) of the population tuning curve was measured for each condition. In the box plots, the central line indicates the median, the box edges indicate the 25th and 75th percentiles, and the whiskers represent the data range. Statistical comparisons between *prior* and *no prior* conditions were made using paired *t*-tests for each contrast. \*\*\**P* < 0.001. **E** Quantification of pursuit bias. The magnitude of bias toward the prior direction was compared between *prior* and *no prior* conditions at each stimulus contrast. Here, bias was defined as the deviation of pursuit direction toward the prior direction relative to the *no prior* condition. Box plot conventions are as described in (D). Paired *t*-test revealed a significantly increased bias under the low-contrast *prior* condition. \*\*\**P* < 0.001. **F** Quantification of directional variability. The standard deviation (SD) of pursuit direction was computed for each condition. Box plot conventions are as described in (D). Paired *t*-tests for each contrast confirmed that directional variability decreased under the low-contrast *prior* condition. \*\*\**P* < 0.001. H/N: high contrast, *no prior*; H/P: high contrast, *prior*; L/N: low contrast, *no prior*; L/P: low contrast, *prior*.

We introduced a direction-dependent decrease in membrane potential, such that neurons whose preferred directions were farther from the prior received stronger inhibitory input, reducing their excitability (Fig. 6A, B; see Methods). We then examined whether this biophysical implementation could reproduce the prior-dependent changes in neural and behavioral responses observed experimentally. In the model, *prior* and *no prior* conditions were defined by the presence or absence of this membrane potential modulation, while high- and low-contrast conditions were determined by the strength of directional input to the network.

Population neural activity showed no significant difference between the *prior* (expecting 0°) and *no prior* conditions when the target, moving toward 0°, was presented as a high-contrast motion stimulus (paired *t*-test, *t*(99) = 0.907, *P* = 0.367; Fig. 6C, D). Under the low-contrast stimulus condition, however, overall population activity was reduced and was further suppressed in the presence of prior expectation compared with the *no prior* condition. Specifically, neurons whose preferred directions differed substantially from the prior direction exhibited the strongest response reduction. This suppression sharpened the population tuning curves, indicating that prior expectation enhances the precision of population coding when sensory evidence is weak (ANOVA, *F*(3, 396) = 166363.934, *P* < 0.001; paired *t*-test, *t*(99) = 221.546, *P* = 2.774×10^-135^; Fig. 6C, D).

The tuned inhibitory modulation also reproduced two key features of prior-driven pursuit behavior. First, pursuit bias was small under the high contrast regardless of prior expectation (Fig. 6E). Under low contrast conditions, however, eye trajectories were significantly biased toward the prior direction (0°) when prior expectation was present (paired *t*-test, *t*(99) = 15.751, *P* = 1.015×10^-28^; Fig. 6E), consistent with the monkeys’ eye movements (Fig. 1D). Second, pursuit variability did not differ between conditions under high contrast (paired *t*-test, *t*(99) = 1.817, *P* = 0.072; Fig. 6F) but was significantly reduced by prior expectation under low contrast (ANOVA, *F*(3, 396) = 10145.923, *P* < 0.001; paired *t*-test, *t*(99) = 56.528, *P* = 3.661 × 10^−77^; Fig. 6F). These results demonstrate that directionally tuned inhibitory modulation of membrane potential is sufficient to account for both the directional bias and the reduction in variability of pursuit behavior induced by prior expectation.

#### Neural evidence of tuned synaptic inhibition observed in low-frequency local field potential activity

Our modeling results suggest that prior expectations modulate sensorimotor behavior and population tuning by adjusting neuronal firing thresholds, implicating tuned synaptic inhibition as the underlying mechanism. To seek empirical evidence for this modulation *in vivo*, we examined baseline pre-stimulus neural activity recorded from area MT in two monkeys performing the smooth pursuit task. Because direct measurement of intracellular membrane potentials in behaving primates remains technically challenging, we utilized the local field potential (LFP) as a proxy for local network excitability and subthreshold synaptic activity^32^.

While classical studies under anesthesia map the phase of low-frequency LFPs to discrete UP and DOWN membrane potential states^33^, low-frequency LFP signals are also known to continuously track network excitability in the awake brain. Okun et al. (2010) provided direct evidence for this link by showing, through simultaneous intracellular and extracellular recordings, that the LFP tracks subthreshold membrane potential fluctuations in unanesthetized animals^34^. Crucially, this relationship has been confirmed in behaving non-human primates: using *in vivo* whole-cell patch-clamp recordings in alert macaques, Tan et al. (2014) demonstrated that spontaneous subthreshold membrane potential excursions during visual fixation are tightly correlated with concurrent LFP deflections^35^. On the basis of these findings, we used the phase and power of low-frequency LFP signals as a continuous index of local network excitability and synaptic input, and examined whether pre-stimulus low-frequency LFP dynamics covaried with prior expectation in a direction-selective manner—serving as an empirical test of the tuned inhibition predicted by our model.

##### (1) Direction-specific inhibition of spontaneous delta phase

We recorded LFPs from area MT using multichannel electrodes, yielding 341 channels from monkey A and 387 channels from monkey B. To test whether low-frequency phase modulation reflected tuned inhibition, we related phase shifts to the angular distance between each channel’s preferred direction and the prior direction. Preferred directions were estimated from an independent direction tuning task (distinct from the main pursuit task; see Methods). We focused on high-gamma activity for this estimation, as it closely reflects local multiunit spiking^36-38^. Direction tuning curves were constructed from the average high-gamma amplitude within 50–300 ms after stimulus onset (Fig. 7A). The distribution of channels as a function of the angular distance between their preferred direction and the prior direction is shown in Fig. 7B.

**Fig 7:**
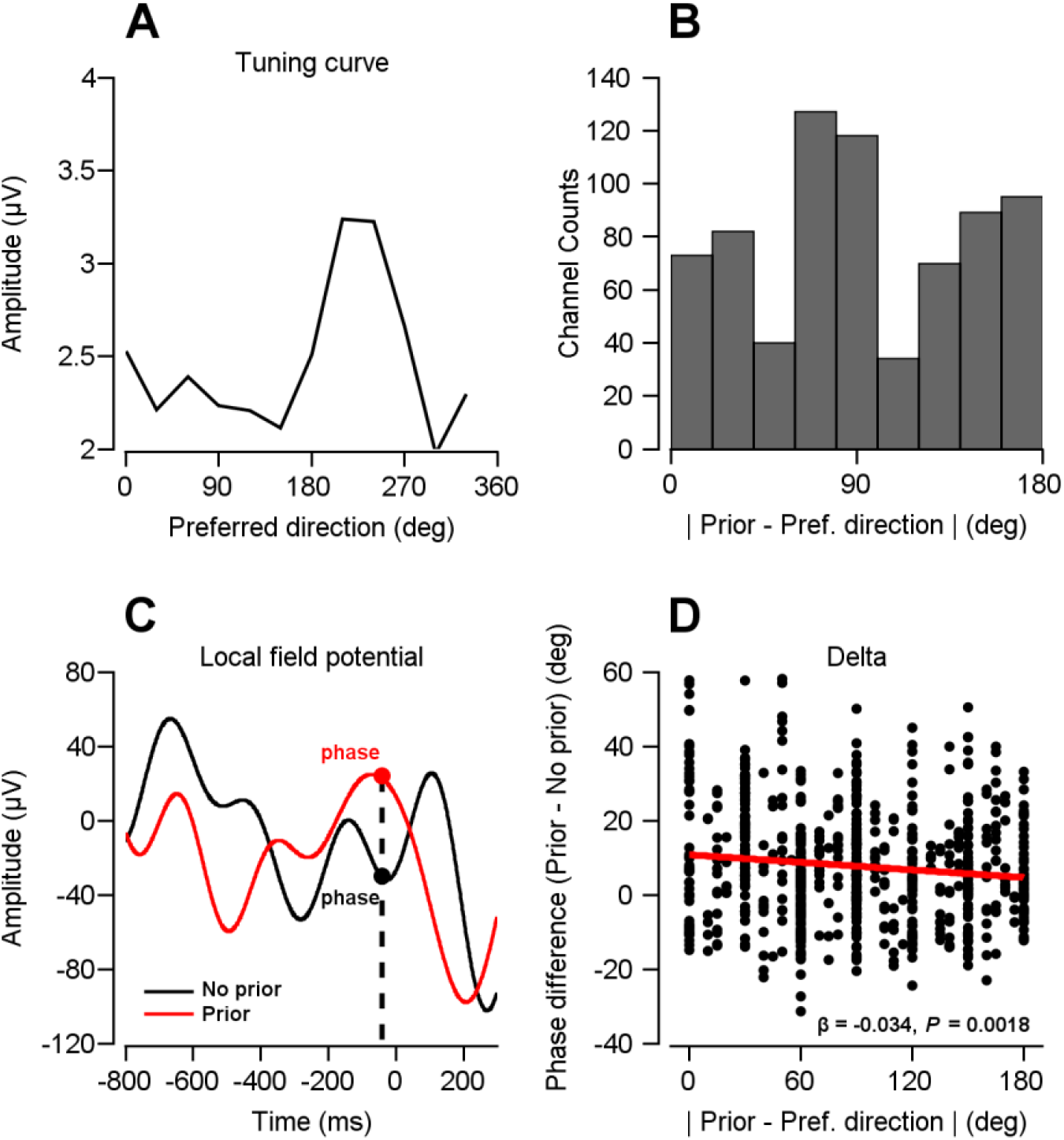
LFP direction tuning in area MT and delta-band phase differences. **A** Representative LFP direction tuning curve recorded in area MT. LFPs were obtained while monkeys performed a direction tuning task. Band-pass filtering (70–150 Hz) was applied to extract high gamma activity and estimate tuning curves of individual recording channels. **B** Distribution of recording channels as a function of the angular difference between the prior direction and each channel’s preferred direction. **C** Schematic of the phase analysis. The example shows delta-band filtered (0.5–4 Hz) LFP signals from a single trial. The black line represents a *no prior* trial, and the red line represents a *prior* trial. Phase values were extracted during the pre-stimulus fixation period. To obtain robust phase estimates, phase values were averaged over the interval from −200 ms to 0 ms relative to stimulus onset. **D** Delta-band phase difference plotted as a function of the angular difference between the prior direction and each channel’s preferred direction (from panel B). Each point represents an individual neuron. The red line indicates a linear regression fit (β is regression coefficient, significance expressed as *P* value).

Using these preferred directions, we examined phase dynamics in the delta frequency range during the pre-stimulus interval as monkeys performed the main pursuit task (Fig. 7C). In the *no prior* condition, delta phase showed no significant dependence on angular distance from the prior direction (regression coefficient = −0.025 at −600 ms, *P* = 0.062; −0.019 at −200 ms, *P* = 0.125; Supplementary figure 8). Under the *prior* condition, however, delta phase decreased systematically with increasing angular distance (regression coefficient = −0.070 at −600 ms, *P* = 5.815 × 10^−5^; −0.051 at −200 ms, *P* = 0.012; Supplementary figure 8). This pattern indicates a structured shift in low-frequency LFP phase as a function of the mismatch between the prior and preferred directions, consistent with tuned modulation of network state during the pre-stimulus period. The decreasing trend was evident both at individual time points (e.g., −600 ms and −200 ms before stimulus onset) and when averaged across the pre-stimulus window (−200 to 0 ms; regression coefficient = −0.034, *P* = 0.002; Supplementary figure 8, Fig. 7D). A marginally significant decrease was also observed in the theta band (regression coefficient = −0.017, *P* = 0.044; Supplementary figure 9A), but no significant tuned modulation was found in other frequency bands (alpha: −0.004, *P* = 0.614; beta: −0.006, *P* = 0.385; low gamma: −0.012, *P* = 0.115; high gamma: −0.010, *P* = 0.160; Supplementary Figure 9B–E).

**Supplementary Figure 8:**
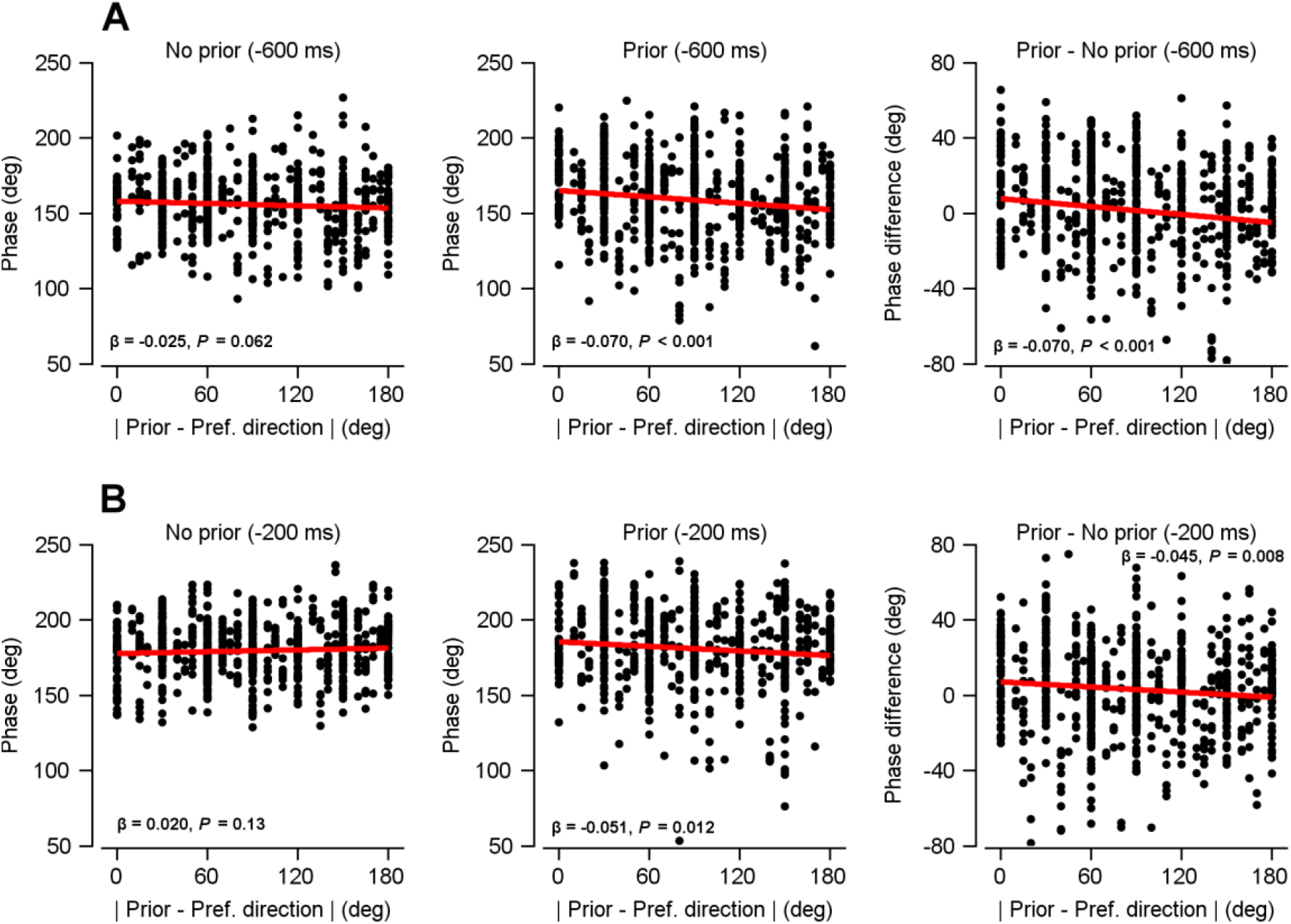
Delta-frequency phase differences in area MT LFP signals between *prior* and *no prior* conditions. **A** For area MT LFP signals bandpass filtered at delta frequency (0.5–4 Hz), the phase at -600 ms during the pre-stimulus fixation interval was used for analysis. (Left) Phase as a function of the angular difference between the prior direction and each channel’s preferred direction in the *no prior* condition. (Middle) Same in the *prior* condition. (Right) Phase difference between the two conditions. Each point represents an individual recording channel. The red line indicates a linear regression fit (β = regression coefficient; significance expressed as *P* value). **B** Phase at -200 ms.

**Supplementary Figure 9:**
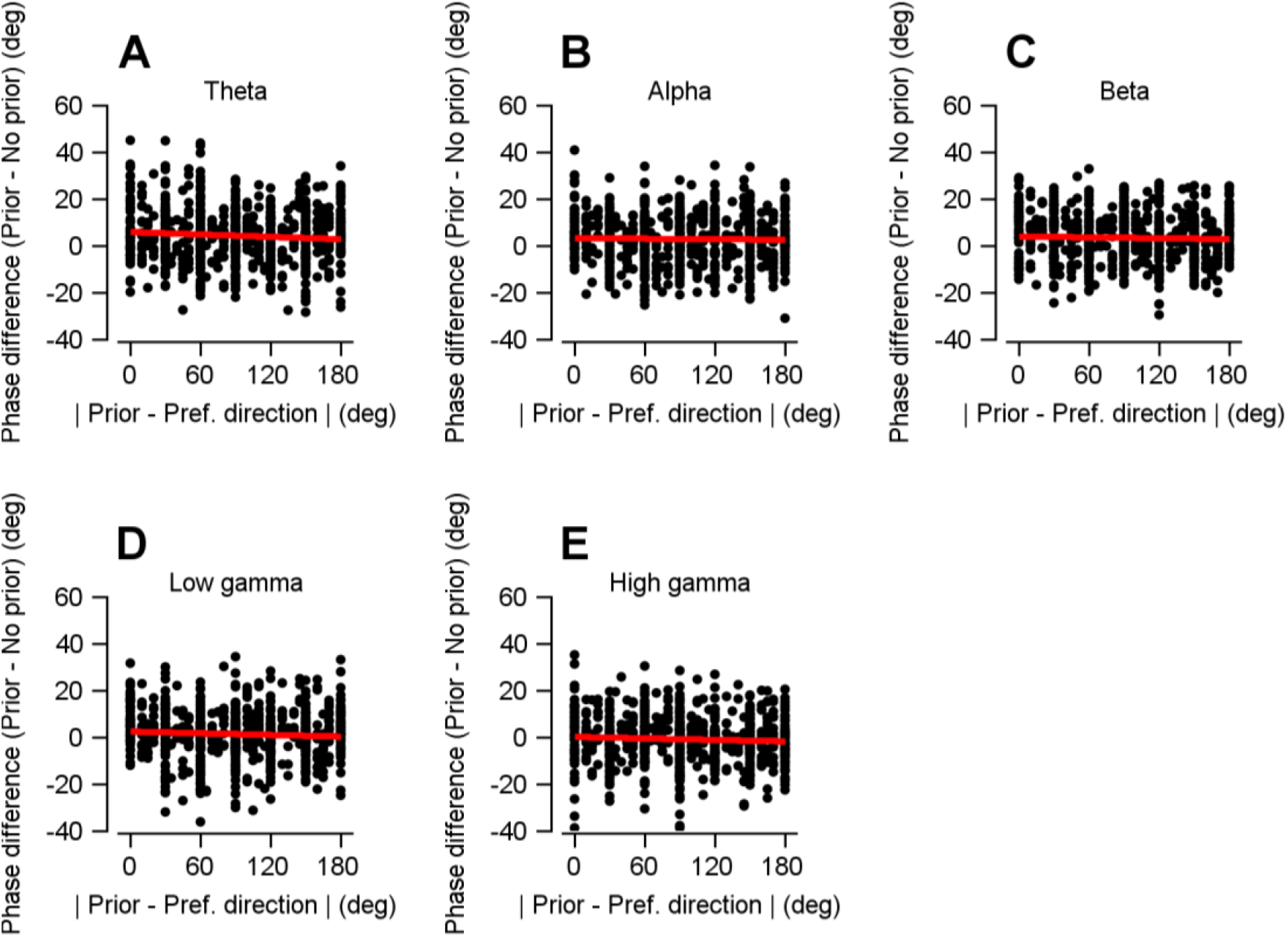
Phase differences across non-delta frequency bands between *prior* and *no prior* conditions. **A** Phase differences in the theta (4–8 Hz) band, plotted as a function of the angular difference between the prior direction and each channel’s preferred direction, as defined in Figure 7B. Each point represents an individual recording channel. The red line indicates a linear regression fit. **B-E** Phase differences in the alpha (8–13 Hz), beta (13–30 Hz), low gamma (30–70 Hz), and high gamma (70–150 Hz) frequency bands.

##### (2) Direction-specific suppression of spontaneous delta power

We next examined whether prior expectation modulated oscillatory power across frequency bands. Compared with the *no prior* condition, the *prior* condition showed a significant decrease in power within the delta, theta, and alpha bands during the pre-stimulus interval (cluster-based permutation test, 10,000 permutations, α = 0.05, 0.5 to 13 Hz; Supplementary figure 10). This low-frequency power suppression was consistently observed in both monkeys.

**Supplementary Figure 10:**
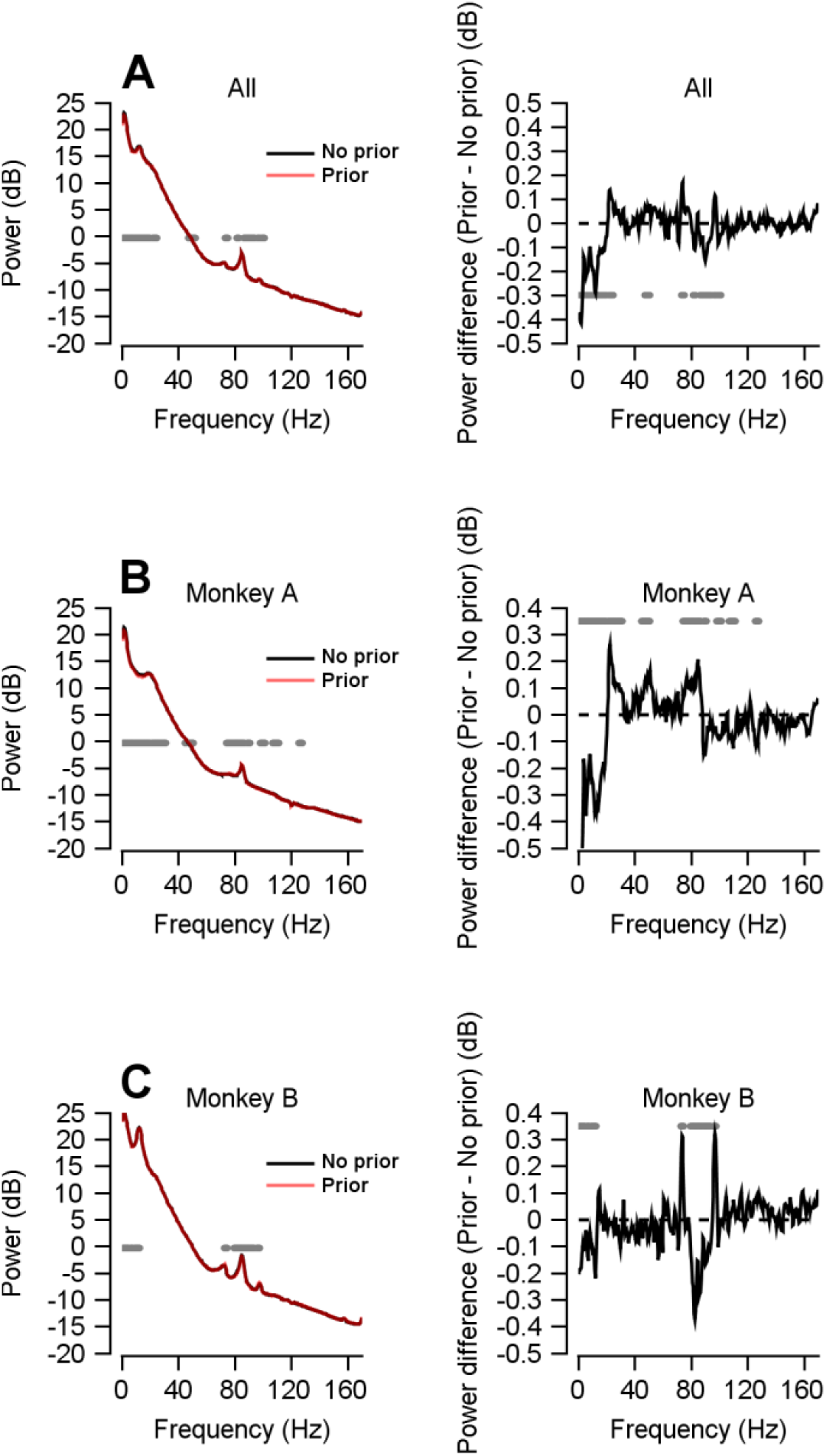
Power spectra for *prior* and *no prior* conditions in two monkeys. **A** (Left) Power as a function of frequency for the *prior* and *no prior* conditions in two monkeys. Spontaneous responses in area MT were analyzed during the fixation period immediately preceding stimulus onset (−800 ms to 0 ms) while the monkeys performed a smooth pursuit eye movement task. The black line represents the *no prior* condition, and the red line represents the *prior* condition. Frequency bands showing significant power differences between conditions were identified using cluster-based permutation tests (10,000 permutations, α = 0.05). (Right) Power difference between the *prior* and *no prior* conditions. **B**–**C** Power spectra and power differences shown separately for monkey A and monkey B.

We next examined whether the low-frequency power reduction observed during the pursuit task (Supplementary figure 10) was tuned to the angular distance between the prior and each channel’s preferred direction. Directionally tuned suppression was found exclusively in the delta band, where power decreased significantly with increasing angular distance (regression coefficient = −2.923 × 10^−3^, *P* = 7.132 × 10^−9^; Supplementary figure 11). No such tuning was observed in any other frequency band (theta: −5.136 × 10^−4^, *P* = 0.202; alpha: −1.489 × 10^−4^, *P* = 0.658; beta: 4.840 × 10^−5^, *P* = 0.841; low gamma: −6.439 × 10^−5^, *P* = 0.777; high gamma: −2.974 × 10^−4^, *P* = 0.296; Supplementary figure 11).

Although a weak trend was present, we did not find significant evidence of tuned inhibition in high-gamma LFP power. This is somewhat surprising given that spontaneous spiking activity at the single-neuron level exhibited signatures of tuned inhibition (Supplementary Figure 6A). This apparent discrepancy may reflect a signal-to-noise limitation: spiking activity provides a more precise measure of neuronal output than high-gamma LFP power, which aggregates signals across a local population. Given the modest effect size in the spike data, it is plausible that the corresponding modulation in high-gamma LFP power was too weak to reach statistical significance.

Taken together, these results demonstrate that prior expectation selectively modulates low-frequency (delta-band) activity in area MT in a direction-dependent manner. Channels whose preferred directions deviate farther from the prior exhibit stronger shifts in low-frequency phase and greater suppression of slow fluctuations. Although these extracellular measures do not provide a direct readout of intracellular membrane potential, the observed LFP modulations are consistent with the hypothesis that prior expectation biases baseline population excitability in a direction-selective manner. These physiological signatures provide empirical support for the threshold modulation mechanism identified by our computational models, linking prior expectation to tuned synaptic inhibition in sensory cortex.

**Supplementary figure 11:**
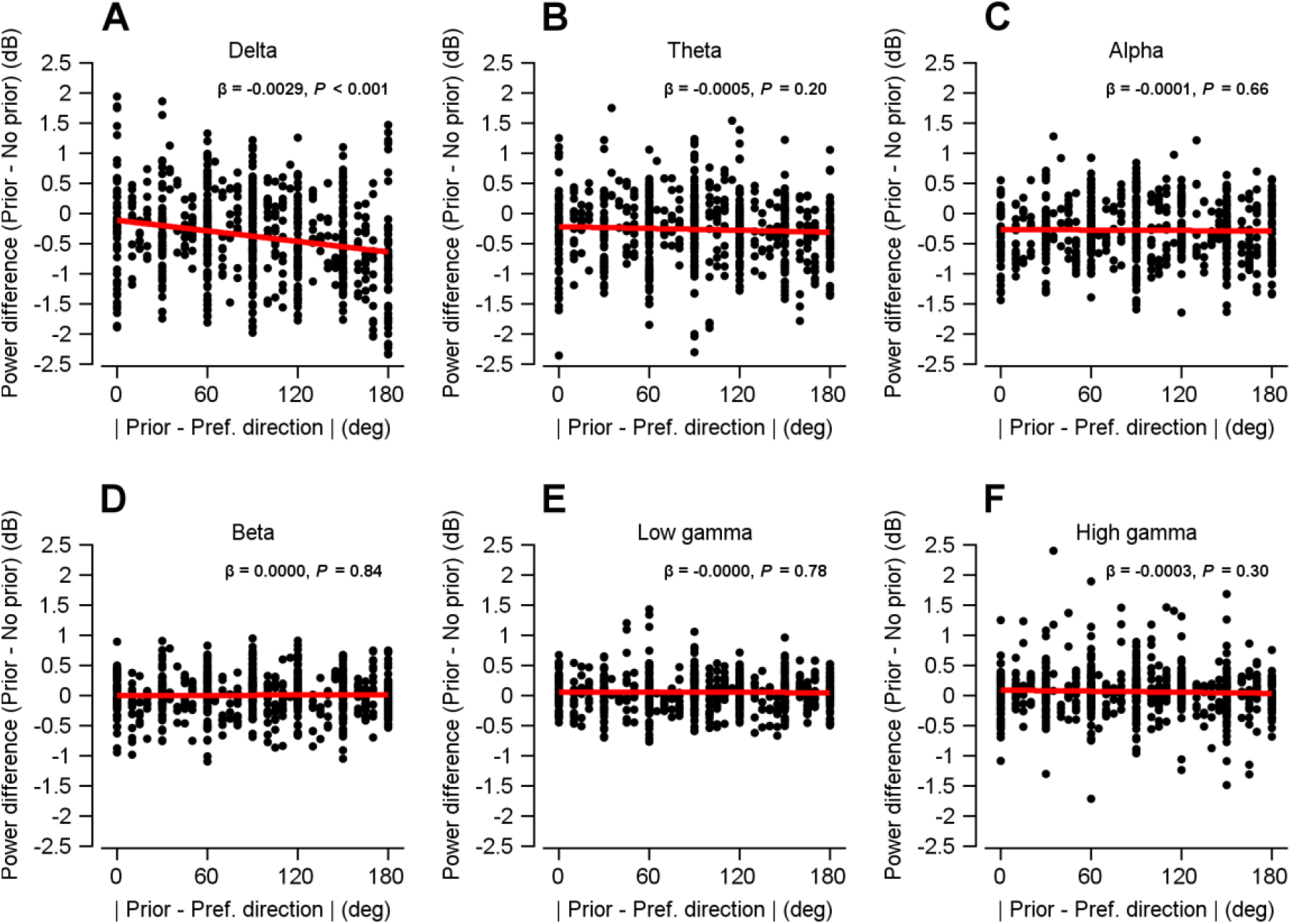
Frequency-specific power differences. **A** Power difference as a function of the angular difference between the prior direction and each channel’s preferred direction, as defined in Figure 7B. For the delta band (0.5–4 Hz), each point represents an individual recording channel. The red line indicates a linear regression fit (*β* denotes the regression coefficient, significance expressed as *P* value). **B**–**E** Power differences in the theta (4–8 Hz), alpha (8–13 Hz), beta (13–30 Hz), low gamma (30–70 Hz), and high gamma (70–150 Hz) frequency bands.

## Discussion

In this study, we investigated how prior expectations influence smooth pursuit eye movements and neural population tuning using monkey behavioral experiments, recurrent neural network (RNN) and biophysical spiking neural network models, and local field potential (LFP) recordings from area MT. Our findings provide converging computational and physiological evidence that prior expectations sharpen population responses and improve behavioral precision under sensory uncertainty, and point to a cellular-level mechanism through which Bayesian inference may be implemented in cortical circuits.

Monkeys’ pursuit eye movements showed systematic biases toward the expected direction, especially under low-contrast conditions where sensory reliability was reduced (Fig. 1D, E). Trial-to-trial variability was also reduced in the presence of priors (Fig. 1F, G), consistent with Bayesian frameworks in which priors exert greater influence when sensory evidence is weak^17,18^. These results align with previous studies showing that expectations improve perceptual estimates of motion, orientation, and object location under ambiguous conditions^9,13,39^.

At the neural level, earlier works have shown that such behavioral effects are accompanied by sharpening of population tuning^3,11,13,17,18^. Consistent with these findings, we observed that prior expectations sharpen direction tuning in area MT by selectively suppressing responses of neurons preferring directions inconsistent with the prior, with the strongest effects under noisy sensory conditions (replicated in Fig. 1H, I).

Such selective suppression is thought to reduce noise in population activity, thereby increasing perceptual accuracy. Importantly, tuned inhibitory mechanisms are well established as a means of sharpening feature selectivity across cortical circuits^40-42^, supporting the plausibility that expectation-related modulation in area MT operates through similar principles. For example, optogenetic activation of parvalbumin-positive interneuron in mouse primary visual cortex sharpens orientation tuning and improves perceptual discrimination^41^. Similarly, in the rat somatosensory cortex, GABA_A_-mediated inhibition preferentially suppressed responses to non-principal whiskers, whereas blocking inhibition broadens tuning^40^. Together, these findings across sensory modalities highlight a general principle whereby structured inhibitory inputs refine population tuning by selectively suppressing non-preferred features.

### Threshold modulation (prediction) vs. Gain modulation (attention)

A key contribution of our modeling is the identification of threshold modulation as the mechanism by which prior expectations influence sensory processing. Attentional modulation is commonly described in terms of gain control: feature attention is typically implemented via input gain, whereas spatial attention is associated with response gain that multiplicatively scales neuronal output^43-45^. However, systematically manipulating either input gain or response gain in our RNNs failed to reproduce the tuning sharpening and variability reduction observed in the behavioral and neural data (Supplementary figure 3, 4).

In contrast, threshold modulation—implemented as elevated firing thresholds for neurons tuned away from the expected direction—captured all major empirical effects. By selectively suppressing weakly driven, feature-irrelevant neurons without globally amplifying activity, this mechanism sharpened population tuning and reduced trial-to-trial variability under low sensory reliability (Fig. 3). These results generalized to a biophysical spiking network, in which threshold modulation was implemented as direction-dependent inhibitory shifts in membrane potential, reproducing the same pattern of tuning sharpening, behavioral bias, and variability reduction (Fig. 6).

An important question raised by these results is why threshold modulation, but not gain modulation, can account for both reduced pursuit variability and sharpened population tuning under low-contrast conditions. Input gain scales sensory signals and response gain scales neuronal outputs; however, both act largely proportionally across the population and therefore preserve the relative tuning structure across neurons. As a result, gain modulation lacks the selectivity required to preferentially suppress neurons whose preferred directions are inconsistent with the prior. In contrast, threshold modulation introduces a nonlinear, selective effect: neurons receiving weak sensory drive—particularly those tuned far from the expected direction—are preferentially silenced, whereas neurons aligned with the prior remain active. This selective suppression sharpens population tuning and reduces variability by limiting contributions from noisy, weakly driven neurons. Because threshold shifts have minimal effects when sensory drive is strong, this mechanism also explains why prior-induced modulation emerges under low sensory reliability but is negligible under high-contrast conditions.

These results indicate that prediction operates through a computational process distinct from attention. Rather than selectively facilitating relevant signals, priors act primarily by inhibiting irrelevant ones through targeted changes in neuronal excitability.

### Flexible tuning through neuronal nonlinearities

A second central issue concerns how the brain flexibly incorporates expectations on short timescales, even on a trial-by-trial basis. While prior information could be implemented through changes in synaptic connectivity, rapidly varying expectations would make this strategy computationally inefficient as a fast control mechanism. Our results instead highlight modulation of neuronal nonlinearities—such as firing thresholds, which largely depend on the overall excitability state of the neuron—as an effective means of flexibly reshaping functional responses without altering core sensory connectivity.

This does not imply that synaptic mechanisms are irrelevant. Rather, prior expectations may be implemented through synaptic and circuit-level processes that manifest functionally as changes in neuronal input-output nonlinearities. Neuronal physiology provides strong support for this view. Whereas AMPA receptor-mediated currents scale approximately linearly with input strength^46,47^, NMDA receptors exhibit pronounced nonlinearity: channels remain largely inactive until membrane depolarization reaches a critical level, at which point Mg^2+^ block is relieved and calcium influx occurs^48,49^. Such threshold-dependent gating enables state-dependent computations without large-scale synaptic rewiring and support flexible operations including memory consolidation, coincidence detection, and flexible state transitions^50-52^.

By analogy, modulation of neuronal input–output nonlinearities in area MT provides a computationally efficient way to reshape population responses according to prior expectations while preserving stable feature selectivity (Fig. 3A). In our RNN analysis, a standard approach—introducing prior information as an additional input and updating synaptic weights across conditions—led to broadened tuning and elevated activity levels that were inconsistent with experimental observations (Fig. 5). In contrast, implementing priors through threshold modulation reproduced both the sharpening of population tuning and the reduction of trial-to-trial variability observed in neural activity and behavior (Fig. 3), with comparable effects obtained in the spiking network model through directionally tuned inhibitory modulation of membrane potential (Fig. 6).

At the biological level, the tuned threshold modulation identified here could be realized through multiple neural substrates, including direction-selective inhibitory circuits, shifts in excitation-inhibition balance mediated by specific interneuron populations, or neuromodulatory influences on baseline excitability^53-55^. Rather than specifying a single cellular mechanism, our results point to a functional principle: direction-specific modulation of neuronal nonlinearity shapes sensory responses under prior expectation.

Finally, the distinction between incorporating contextual information through synaptic weight-based integration and through modulation of neuronal input-output nonlinearities may reflect broader architectural differences across cortical hierarchies. Higher-order areas such as the prefrontal cortex (PFC) often exhibit mixed selectivity and integrate contextual variables directly alongside sensory inputs^15,26^. For example, Mante et al. (2013) modeled sensory and contextual signals as inputs to a recurrent neural network, capturing how PFC neurons flexibly combine sensory and cognitive variables^26^. In contrast, sensory areas such as area MT maintain stable, feature-specific representations and may incorporate contextual information through modulatory signals that reshape response nonlinearities rather than altering core sensory connectivity. From this perspective, modulation of neuronal nonlinearity offers a biologically plausible and computationally efficient strategy for incorporating prior expectations into sensory processing.

An important clarification concerns the scope of our modeling approach. The RNN was not trained to acquire prior expectations from experience; rather, it was first trained under the *no prior* condition to establish a baseline sensorimotor mapping, and prior effects were subsequently introduced as structured modulations of unit input–output nonlinearities through parameter optimization. The model therefore addresses how an already-formed prior can be implemented within a neural population, not how such priors are learned.

Within this framework, direction-specific modulation of firing thresholds provides a parsimonious mechanism capable of reproducing key empirical observations, including tuning sharpening and reduced trial-to-trial variability under low sensory reliability. However, this should be interpreted as a functional account rather than a specific biological implementation; multiple circuit-level mechanisms—including learned feedback projections, recurrent inhibitory connectivity, or distributed modulatory signals—could give rise to similar effects. Future work will be needed to bridge this functional description with models of how priors are acquired, updated, and represented across cortical circuits.

### Low-frequency top-down modulation

A third major theme concerns the source of the modulatory signal implementing threshold-like inhibition in area MT. Although local synaptic circuity can support rapid sensory encoding and precise feature selectivity (Fig. 2), dynamically updating prior expectations solely through synaptic reconfiguration within area MT would be inefficient given the rapid timescale of expectation changes. Instead, our behavioral, physiological, and modeling results point to low-frequency top-down signals from higher cortical regions—such as PFC, frontal eye field smooth eye movement region (FEFsem), or supplementary eye field (SEF)—as a source of direction-selective modulation that adjusts neuronal excitability in area MT.

Consistent with this interpretation, prior expectations reduced pre-stimulus delta-band power and induced direction-specific phase modulation in area MT (Fig. 7, Supplementary figure 11). Delta-band oscillations have been strongly linked to top-down predictive signaling and are known to convey information from frontal regions to sensory cortices^56,57^. These observations align with hierarchical communication frameworks in which bottom-up sensory signals are carried by higher-frequency activity, such as gamma^58-60^, whereas top-down contextual signals, including priors, are transmitted through low-frequency bands^61-63^. Within this framework, delta-band modulation in area MT likely reflects top-down influences that selectively alter neuronal excitability, preferentially suppressing responses that deviate from the expected motion direction. Notably, this pattern closely matches the direction-specific threshold modulation predicted by our RNN model (Fig. 4), and its biophysical implementation as tuned inhibitory modulation in the spiking network (Fig. 6).

These findings motivate a computational architecture in which prior information is encoded in higher-order regions and transmitted to sensory cortex as frequency specific modulatory signals. Multi-region RNNs comprising a cognitive module that represents prior information and a sensory module that processes incoming inputs provide a promising framework^64,65^. If top-down influences are inhibitory, training such networks on tasks requiring integration of priors with sensory evidence should naturally give rise to structured inhibitory projections from cognitive to sensory modules.

Complementing these modeling approaches, future experiments combining simultaneous recordings from frontal regions that convey expectation signals and from area MT will allow direct tests of these predictions. Such recordings will enable precise characterization of the timing, directionality, and frequency specificity of top-down signals during prediction-driven sensorimotor processing^66,67^.

## Conclusion

In summary, this study demonstrates that prior expectations enhance motor behavior under sensory uncertainty through a threshold-based inhibitory mechanism. Our findings suggest that prior information can be incorporated through rapid and reversible modulation of neuronal input-output nonlinearities, potentially mediated by low-frequency top-down signals from higher cortical regions. This form of modulation sharpens sensory representations, reduces internal variability, and biases behavior toward expected features, offering a biologically grounded implementation of Bayesian inference in sensory–motor systems.

## Methods

### Animals

Two 10-year-old adult male rhesus monkeys (*Macaca mulatta*) weighing 9 to 11 kg were used for the neurophysiological experiments. All research protocols were approved by the Sungkyunkwan University Institutional Animal Care and Use Committee (SKKUIACUC2017-06-10-1). Prior to the experiments, we performed two separate surgeries. A head holder was implanted on the skull for the head restraint, after which a cylindrical chamber fabricated from Poly Ether Ether Ketone (PEEK) was implanted on the skull close to the lunate sulcus for an angled approach to area MT. During the surgeries, each monkey was under isoflurane anesthesia, while antibiotics and analgesics were administered post-operatively to minimize infection and pain. Behavioral and neural data reported in our previous work were reanalyzed in the present study to address new mechanistic questions^13^. Importantly, the local field potential (LFP) results presented here have not been reported previously.

### Task design

Visual stimuli were presented on a gamma-corrected 24″ Cathode-Ray Tube (CRT) monitor (HP1230, 1600 × 1200 pixels, 85-Hz vertical refresh rate). The monitor was placed 570 mm from the animal, subtending 38.67° by 29.49° of the horizontal and vertical visual field, respectively. The background on which the visual stimuli were presented was gray with a luminance level of 36.8 cd/m^2^ (luminance range was 0 to 73.68 cd/m^2^). The presentation of the visual stimuli and recording of the eye movement data were controlled using a real-time data acquisition system (Maestro version 3.3.11). A custom-built photodiode system was used to ensure the accurate timing of the visual stimuli.

The two monkeys were trained in a smooth pursuit eye movement task (Fig. 1A). The pursuit target was a random dot kinematogram, and its size in each day’s experiment was determined as one of 4° × 4°, 8° × 8°, or 12° × 12° depending on the location and size of the receptive fields of the recorded MT area. Each trial began when the animals fixated their eyes on a small dot (fixation spot) at the center of the monitor screen. After a randomized fixation duration (800, 1300, or 1800 ms), the dot patch appeared at the center of the screen or 1 to 2° displaced from the center to the opposite direction of the target direction. This positional manipulation was used to facilitate the initiation of smooth pursuit and did not qualitatively affect the measured behavioral outcomes; data from different initial positions were pooled for analysis. Subsequently, a local motion, whereby all the dots inside the invisible circular window moved in the target direction at a given speed but the invisible window did not move, was implemented. Following the local motion, both the dots and windows moved together at the same speed and in the same direction as the local motion for 500 to 700 ms. Behavioral analyses and RNN training were based on eye velocity measured during the open-loop period (150 ms after pursuit initiation^68^), to exclude the influence of the feedback signals. This analysis window typically spans the late phase of local motion and the early phase of global motion, thereby capturing pursuit responses driven by the initial motion signal. If the animals maintained their eyes at the center of the pursuit target within a 4° window during the target movements, then they were rewarded with a drop of water or juice. The precision of the sensory evidence (sensory stimulus) was controlled by randomly selecting a pursuit target stimulus from the two stimulus types in each trial: One was a random-dot kinematogram with 100% luminance contrast (high contrast), and the other was a random-dot kinematogram with 12 or 8% (on average 10%) (for monkeys A and B, respectively) luminance contrast (low contrast) (Fig. 1B).

To manipulate the animals’ prior expectation of the motion, we used two block types: *prior* and *no prior* blocks (Fig. 1C). In the *no prior* block, the fixation spot was red, and the pursuit target direction was randomly and evenly selected from three directions spaced 120° apart, resulting in broadly distributed directional expectations. In the *prior* block, the fixation spot was green, and the target moved in one of the three narrowly spaced directions (a central direction and ±15°), with the central direction presented twice as often as the other two directions. This design encouraged the animals to develop a strong expectation for the central direction, which we termed prior direction. The prior direction selected based on the preferred directions of the recorded neurons.

The distribution of directions in the prior block was designed to simultaneously manipulate both the probability and the geometric spacing of stimuli to maximize the strength of the induced expectation. By increasing the probability of the central direction while clustering the possible directions within a narrow range, the task encouraged the formation of a strong behaviorally relevant expectation under sensory uncertainty. This design choice does not isolate probability-based priors from factors such as central tendency or geometric structure; rather, it intentionally combines them to amplify expectation-related modulation in both behavior and neural activity.

Importantly, the prior directions was included in both blocks, enabling direct comparison of behavioral and neural responses to identical sensory stimuli under different expectation contexts. The *prior* and *no prior* blocks consisted of 252 and 378 trials, respectively. The first block of each session was randomly selected, and the two block types alternated thereafter. We typically collected four blocks per condition, yielding approximately 2,520 trials in total.

### Behavior analysis

The horizontal and vertical eye positions were separately recorded at a sampling rate of 1 kHz using an infrared video tracking system (EyeLink 1000 Plus, SR Research Ltd.). A low-pass Butterworth filter with an order of two and a cutoff frequency of 20 Hz was applied on the horizontal and vertical components of the eye position. Subsequently, the filtered eye positions were differentiated to obtain the horizontal and vertical eye velocities. To focus on the initiation of the smooth pursuit eye movements and remove the influence of the saccadic eye movements (saccades) on the behavioral responses, the trials with saccades occurring within a time window between −100 and 250 ms from the onset of the pursuit target were removed. Next, the eye velocity traces were rotated such that their average direction was 45°. The rotated eye velocity during the open-loop period [150 ms from pursuit onset^68^] was decomposed by horizontal and vertical templates^69^, which were the average of each rotated eye velocity component between −20 and 100 ms from the average pursuit latency (mean pursuit onset for high- and low-contrast stimuli: 76 and 123 ms in monkey A and 57 and 105 ms in monkey B). The optimal pursuit latency and scaling factors were estimated through the sliding and scaling of the two templates using the least-squares method and the nonlinear optimization with mesh adaptive direct search (NOMAD) algorithm^70^. We included trials for further analysis only if the fitted function accounted for more than 70% of the data variance. After this procedure was carried out, we used the results for subsequent analysis only if the number of remaining trials of the prior direction in each block for the day’s experiment was more than 70. The eye velocities were normalized by dividing them by the target speed used in the motion stimulus. This was possible because there was no significant difference in speed between the *prior* and *no prior* conditions^13^. This normalization allows direction to be analyzed independently. Importantly, it enables the use of a large number of eye velocity datasets for training RNN models, thereby improving the efficiency and speed of the training process.

### Neural population direction tuning curves

Neural recordings were obtained from area MT and used to construct directional population tuning curves. Detailed experimental procedures for neural recording, spike sorting, and receptive field characterization have been reported previously^13^ and are summarized briefly here. Extracellular activity was recorded from well-isolated area MT neurons while monkeys performed the smooth pursuit task. Only neurons with receptive fields near the fovea were included, as pursuit targets were presented at or near the fovea. Spike sorting was performed offline using principal component analysis and waveform features (Offline Sorter, Plexon). Only neurons with clearly separable spike clusters and reliable stimulus-evoked responses were included in the analysis. Across recording sessions, we obtained responses from 137 neurons in monkey A and 120 neurons in monkey B.

To quantify neural responses during the pursuit task, peristimulus time histograms (PSTHs) were constructed for each neuron and condition, and smoothed using a Gaussian kernel with a standard deviation of 10 ms. Spike counts were measured within a 100-ms time window aligned to each neuron’s response latency, which was estimated from the mean PSTH for each contrast condition (high or low). Neurons without clearly identifiable stimulus-evoked responses were excluded. This yielded datasets of 136 and 111 neurons for high- and low-contrast conditions in monkey A, and 117 and 110 neurons in monkey B, respectively.

The preferred motion direction of each neuron was determined in a separate direction tuning task performed before the smooth pursuit experiment. During the pursuit task, neural responses were analyzed across the four experimental conditions defined by stimulus contrast (high *vs*. low) and prior expectation (*prior vs. no prior*). Population direction tuning curves were constructed by aligning individual neuronal responses according to their preferred directions for each condition. This procedure allowed us to assess how prior expectations modulated the sharpness of population tuning under different levels of sensory reliability.

### RNN model

#### Construction of the network architecture

To investigate a system in which eye velocity profiles of the smooth pursuit eye movement task could be read out over time, we implemented the dynamical system (Fig. 2A), using a standard continuous RNN equation of the form

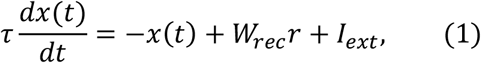

where *τ* corresponds to the synaptic decay time constants for the *N* units in the network, *x* is the synaptic current variable, *W*_*rec*_ is the recurrent connectivity matrix, *r* is the firing rates in the network, which were related to the activations by the rectified hyperbolic tangent function (input-output function or activation function), such that 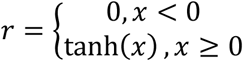. For all simulations, we used a fixed time constant of 30 ms for *τ*, and *N* was fixed at 250.

The elements of *W*_*rec*_ was initialized as a random matrix from a normal distribution with zero mean and an SD of 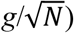, where the synaptic scaling factor, *g*, was set at 1.5^19,71,72^.

The external currents (*I*_*ext*_) included task-specific input stimulus signals:

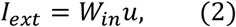

where the time-varying stimulus signals (*u*) was fed to the network through the input weight matrix *W*_*in*_, which was initialized to have zero mean (normally distributed values with 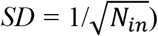^19,71,72^. *N*_*in*_ corresponds to the number of input signals. The input to the neural network represented the direction vector of target velocity. Directional information was encoded using a two-dimensional vector (*cos(θ), sin(θ)*), where *θ* denotes the target motion direction. To account for inter-trial directional variability, Gaussian noise with a standard deviation of 7°, matching the average variability observed in monkey behavior^13^, was added to *θ* on each trial. Sensory contrast was implemented by scaling the overall amplitude of the input pulse while preserving the directional ratio defined by the *cos(θ), sin(θ)*. Specifically, the input pulse amplitude was set to 1 for the high-contrast condition and 0.1 for the low-contrast condition, corresponding to the 100% and 10% contrast stimuli used in the smooth pursuit task. Thus, contrast manipulation altered the strength of the sensory input without changing the encoded motion direction. To examine spontaneous activity in the trained neural network, the input is as follows.

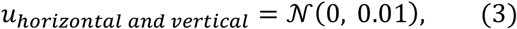

where 𝒩(0, 0.01) represents a Gaussian random noise with zero mean and variance of 0.01. Eq.1 is discretized using the first-order Euler approximation method:

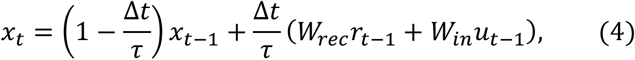

where Δ*t* = 1 ms is the discretization time step size used throughout this study.

To produce the two desired velocity outputs, which were the horizontal- and the vertical-velocity of the smooth pursuit eye movement task, we defined a linear readout of the population activity:

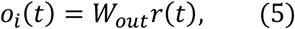

where *o* represents the two velocity readouts (*i* = 1,2) and is a linear combination of the internal firing rates using the readout matrix *W*_*out*_, which was initialized to all zero values.

#### Training the network

Networks were optimized to generate smooth pursuit eye movement. The loss function (ℒ) used to optimize the network considered the difference between the network output (*o*) and the desired eye movement (*z*) which was derived from monkey behavior. Throughout the study, we used the root mean squared error defined as

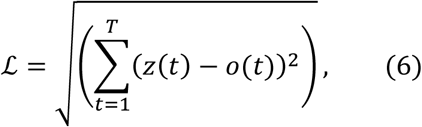

where *T* is the total number of time points in a single trial.

In order to train the neural network to minimize the above loss function, we used the adaptive moment estimation stochastic gradient descent algorithm, known as backpropagation through time (BPTT). The learning rate was set to 0.0001. The gradient descent method was used to optimize the following parameters in the RNN model: input weight (*W*_*in*_), recurrent connectivity weight (*W*_*rec*_), readout weight (*W*_*out*_).

#### Estimation of the preferred direction of neurons

High contrast stimuli were presented to the trained neural network at 1° increments, ranging from -179° to 180°, to obtain the direction tuning curve and corresponding firing rate of each neuron. For each neuron, this procedure was repeated 100 times, and the responses were averaged to generate a representative tuning curve. The preferred direction of each neuron was then estimated by fitting the averaged tuning curve with a linear regression model^73,74^:

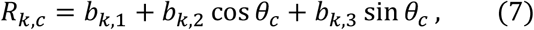

where *R* is the average firing rate of neuron *k* over time during open-loop period [150 ms after pursuit onset^68^] for motion condition *c, b*_*k*,1_, *b*_*k*,2_, and *b*_*k*,3_ are coefficients for offset, cosine and sine, and *θc* is the angle of the current target. A normalized coefficient vector *C*_*k*_ was then defined as:

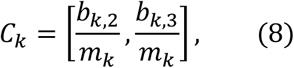

where

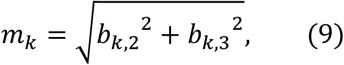

Finally, the preferred direction of each neuron was computed as the angle of *C*_*k*_ using the arctangent function and expressed in degrees:

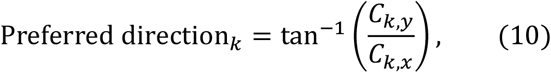

#### Population direction tuning curve

To construct the population direction tuning curve, we first collected the firing rates of individual neurons in response to a specific motion stimulus. The firing rates of each neuron were plotted as a function of their preferred direction. For example, when a stimulus was presented at 0 degrees, the firing rate of each neuron was plotted according to its preferred direction. Neurons with a preference for the 0-degree direction exhibited the highest firing rates, while those with preferred directions farther from 0 degrees showed progressively weaker responses. This approach typically results in a bell-shaped tuning curve, as confirmed by the RNN results in this study (Fig. 2H). To obtain a representative population direction tuning curve, the data were fitted with a Gaussian function.

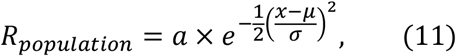

where *a* is maximum firing rate of the gaussian fit, *u* is the center direction, and *σ* is the standard deviation. The sharpness of the population tuning curve was quantified by measuring the full width at half maximum (FWHM), a standard metric for characterizing the width of the tuning curve.

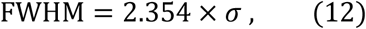

where *σ* is standard deviation estimated from the population direction tuning curve.

#### Neuron-specific modulation of the input-output function parameters

We trained the neural network in two distinct steps. This two-stage procedure was essential for dissociating the learning of baseline sensory–motor dynamics from the modulatory effects of prior expectation on the neurons’ input–output functions. Because prior knowledge is thought to influence sensory processing through network-level inhibitory modulation that flexibly regulates neuronal responses—rather than through changes to fixed connectivity—we first allowed the network to acquire stable sensorimotor dynamics under purely sensory-driven conditions, and only then introduced neuron-specific modulatory changes. This separation ensured that prior-induced effects could be attributed to targeted modulation of input-output function parameters rather than confounding changes in the recurrent circuitry itself.

##### (1) Training the neural network to perform the normal pursuit task

To mimic realistic eye movements under normal conditions, we first trained the neural network using monkey eye movement traces obtained under high and low contrast stimulus conditions, without prior expectation. During this phase, all input-output function parameters (i.e., threshold and gains) were set to 0 and 1. The network, including the input weight (*W*_*in*_), recurrent connectivity weight (*W*_*rec*_), and readout weight (*W*_*out*_), was trained to reproduce smooth pursuit eye movements. Most of the training procedures followed standard methods described above (Fig. 2A). The network was trained for 4,000 iterations (Supplementary figure 12).

##### (2) Fitting the input-output function parameters to account for prior-induced behavior

To enable more precise control of neuronal activity driven by prior expectation, we implemented neuron-specific modulation by independently adjusting the threshold or gain parameters of each neuron’s input-output function (Supplementary figure 2A). In other words, prior-induced neuromodulation was achieved by directly controlling the input–output transformation of each neuron. This approach was motivated by previous studies of motor control showing that modulatory signals can alter neuronal responses by adjusting input–output functions^21^. Following this logic, we hypothesized that prior expectations could be implemented through modulatory changes at the level of neuronal nonlinearity. We obtained the optimal threshold or gain values using backpropagation through time, targeting the generation of eye movement modulated by prior expectation. During this phase, the network architecture - comprising input weight (*W*_*in*_), recurrent connectivity weight (*W*_*rec*_), and readout weight (*W*_*out*_) - remained fixed. The learning process acted solely on the modulatory pathway, iteratively adjusting each neuron’s threshold or gain. To emulate realistic eye movement affected by prior expectation, the input-output function parameters were trained using only averaged prior-induced eye movement biases under high and low contrast stimulus conditions. Note that we did not use the full set of trial-by-trial pursuit traces or variability. After an additional ∼2000 training iterations beyond the initial weight training, the network successfully reproduced prior-induced pursuit bias (Supplementary figure 12).

Again, the input-output (or activation) function *f*, which transforms neuronal activity *x* into firing rates *r*, is defined as:

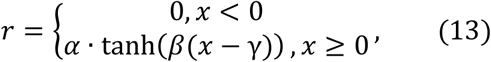

The response gain *α* is the amplitude of the function’s output and thus controls the maximum firing rate of neuron *i* (Supplementary figure 2D). When untrained, its default value is set to 1. The input gain *β* is the function’s slope and thus controls the input-output sensitivity of neuron *i* (Supplementary figure 2C). Its default value is 1 when untrained. The threshold *γ* is the horizontal shift (x-axis) of the function *f* and thus controls the input threshold for neuronal activation (Supplementary figure 2B). Its default value is set to 0 when untrained.

**Supplementary Figure 12:**
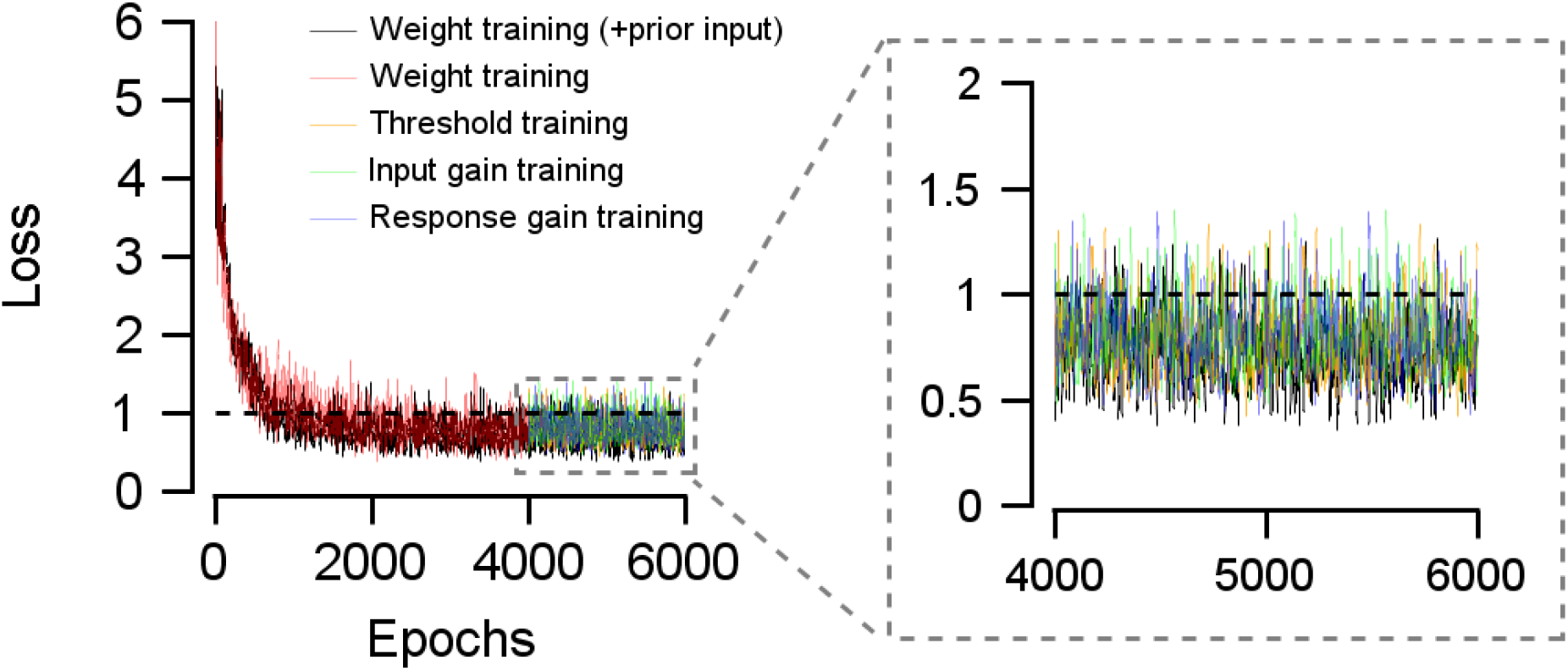
Loss function. The curves show the error between the model’s predicted pursuit behavior and the actual monkey behavior across training iterations. Each line represents a different training method used in this study: black, weight-based optimization performed with prior input (6000 training iterations); red, weight-based optimization performed without prior input (4000 training iterations); yellow, green, and blue, optimization of the threshold, input gain, and response gain parameters, respectively. For the input-output function parameter training (yellow, green, blue), the network was first trained for 4000 iterations using the weight-based optimization without prior input (red). Only the modulatory parameters were then optimized for an additional 2000 iterations to fit the prior-induced pursuit biases. The inset shows an enlarged comparison of the learning outcomes for the weight-based optimization with prior input and the optimization of input-output function parameters. As illustrated, the final loss values were nearly identical across all four methods.

#### Introducing prior input to the neural network

To validate the effectiveness of the modulatory parameter, we conducted a control experiment in which prior information was incorporated directly into the network input instead of altering the modulatory parameter. The input in this control experiment included both the amplitude modulated direction vector of target velocity and the prior information (Fig. 5A). The prior input strength was set to 0.1, such that the *prior* condition had a strength of 0.1, while the *no prior* condition had a strength of 0.

Behavioral testing indicated that excessively strong prior input (e.g., 1) produced effects such as pursuit bias and reduced pursuit variability, not only under low contrast stimuli but also high contrast stimuli, which is inconsistent with actual monkey behavior. Conversely, excessively weak prior input (e.g., 0.01) did not produce significant behavioral changes under either high- or low-contrast condition. Therefore, the prior input strength capable of reproducing the observed behavioral changes in monkeys was selected.

Although the prior input was designed to induce changes in the monkey’s behavior, its effect on neural population activity remained unclear. Importantly, introducing prior information affects network dynamics through synaptic weight updates during training. Therefore, the primary goal of this control experiment was to compare neural population tuning resulting from weight optimization driven by prior input with that arising from direct modulation of neuronal input-output nonlinearities. For the network output, actual monkey eye movement data corresponding to each condition (high contrast, low contrast, *prior*, and *no prior* conditions) were used. The *prior* condition training set included pursuit bias data; thus, it was expected that the RNN would exhibit pursuit bias after training. However, any changes in pursuit variability or neural population tuning beyond this bias can be considered emergent phenomena that are not directly predictable. The learning objective was to optimize the input weights (*W*_*in*_), recurrent weights (*W*_*rec*_), and readout weights (*W*_*out*_) without adjusting the modulatory parameter. The network was trained for 6,000 iterations (Supplementary figure 12). The loss function converged early and reached a stable plateau, indicating that additional training resulted in no meaningful improvement.

### Spiking neural network model

#### Network architecture

To investigate the emergence of stimulus-dependent population activity in a structured recurrent circuit, we implemented a network of *N* = 500 leaky integrate-and-fire (LIF) neurons arranged on a one-dimensional circular feature space representing preferred stimulus direction (Fig. 6A). Each neuron *i* was assigned a preferred angle *ϕ*_*i*_ ∈ [−*π, π*], uniformly spaced across the ring.

#### Neuron dynamics

The membrane potential *V*_*i*_ (*t*) of each neuron evolved according to the standard LIF dynamics:

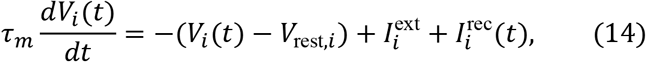

where *τ*_*m*_ = 30 ms is the membrane time constant, *V*_rest,*i*_ is the resting potential, 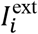 is the external input current, and 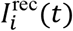 is the recurrent synaptic input. The baseline resting potential was set to *V*_rest_ = −70 mV, and the spike threshold was *V*_th_ = −50 mV. When *V*_*i*_ (*t*) reached threshold, a spike was emitted and the membrane potential was reset to *V*_reset_ = −70 mV.

The model was simulated using a first-order Euler discretization with a time step of Δ*t* = 1 ms:

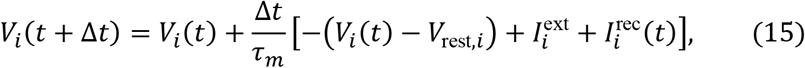

Simulations were run for *T* = 150 ms with a time step of 1 ms. At each time step, synaptic variables were updated, recurrent inputs were computed, and membrane potentials were integrated. Spike events were recorded as binary variables.

#### External input

Each neuron received a directionally tuned external input:

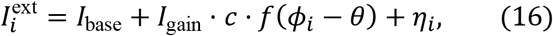

where *I*_base_ = 25 is a constant baseline input, *I*_gain_ = 20 scales the stimulus-dependent component, *c* denotes stimulus contrast, and *θ* is the stimulus direction. The tuning function was defined as:

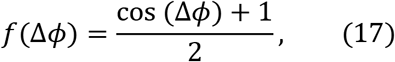

which ensures non-negative input. The noise term *η*_*i*_ was drawn from a Gaussian distribution with standard deviation *σ* = 1.0. Two contrast levels were used in the simulations: high-contrast (*c* = 1.0) and low-contrast (*c* = 0.1).

#### Recurrent connectivity

Recurrent interactions were defined by a distance-dependent “Mexican hat” connectivity profile on the circular manifold. The synaptic weight between neurons *i* and *j* was given by:

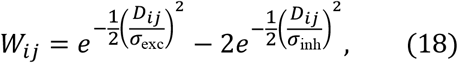

where *D*_*ij*_ denotes the circular distance between preferred angles *ϕ*_*i*_ and *ϕ*_*j*_. The widths of excitation and inhibition were set to *σ*_exc_ = 0.5 and *σ*_inh_ = 2.0, respectively. The weight matrix was normalized by 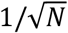 to ensure stable dynamics.

The recurrent synaptic current was computed as:

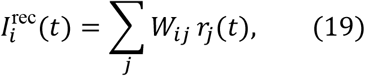

where *r*_*j*_(*t*) represents the synaptic activation variable of neuron *j*.

#### Synaptic dynamics

Spike trains were filtered through an exponential synaptic kernel to generate the synaptic activation variable:

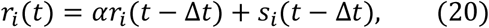

where *s*_*i*_(*t*) is the binary spike train and *α* = exp (−Δ*t*/*τ*_*m*_).

#### Tuned baseline inhibition

To introduce a structured bias in network excitability, we incorporated a direction-dependent modulation of the resting potential (Fig. 6B). Specifically, the effective resting potential of each neuron was adjusted as:

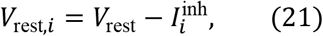

where the inhibition profile increased linearly with the circular distance from the prior direction (0^∘^):

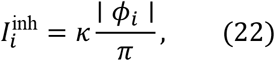

with *κ* = 0.5 controlling the strength of tuned inhibition. This manipulation selectively reduced excitability for neurons tuned away from the prior direction.

### Population vector decoding

To estimate the stimulus direction represented by the population activity, we applied a population vector decoding method to the spiking output of the network. For each trial, firing rates were computed from the spike trains generated by the LIF neurons by averaging spike counts over the simulation window:

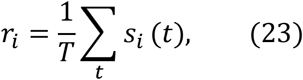

where *s*_*i*_(*t*) denotes the binary spike train of neuron *i* produced by the spiking network model, and *T* = 150 ms is the total simulation duration.

The decoded direction 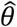 was obtained as the angle of the population vector:

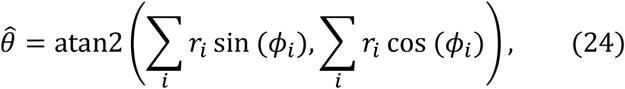

where *ϕ*_*i*_ is the preferred direction of neuron *i*. This method provides a circular estimate of the stimulus direction encoded in the population spiking activity, based on the weighted contributions of all neurons.

### Electrophysiological LFP recordings and LFP power analysis

To examine whether prior expectation modulates baseline neural activity, we compared spontaneous local field potential (LFP) signals between *prior* and *no prior* conditions. LFP signals were analyzed during the baseline period from -800 ms to stimulus onset.

An eight-channel probe (Multi-electrode Type I, Thomas RECORDING) with impedances from 0.5 to 1 megohm (at 1 kHz) was used to record the local field potentials (LFPs) in the middle temporal (MT) area using the Motorized Electrode Manipulator System (Thomas RECORDING). Extracellular electrical signals from the electrode were low pass–filtered at a cutoff frequency of 170 Hz, digitized at a sampling rate of 1 kHz for LFPs (OmniPlex). In total, 341 channels were recorded over 57 days in monkey A, and 387 channels over 64 days in monkey B. LFP signals were recorded with hardware-based line-noise suppression that removed the 60 Hz component. Residual line-noise harmonics were subsequently attenuated using zero-phase finite impulse response (FIR) filters implemented in EEGLAB, specifically via the *cleanline* plugin^75^. Spectral analyses were performed using the Chronux MATLAB toolbox^76^. A Hanning window was applied, and zero-padding was used to interpolate frequency values. Power spectra were computed for the fixation interval, defined as -800 ms to 0 ms relative to stimulus onset.

### LFP phase analysis

The LFP signals were bandpass filtered into six canonical frequency bands—delta (0.5–4 Hz), theta (4– 8 Hz), alpha (8–13 Hz), beta (13–30 Hz), low Gamma (30–70 Hz), and high Gamma (70–150 Hz)— using Hamming-windowed, sinc-based FIR filters implemented in EEGLAB, specifically via the *pop_eegfiltnew* plugin^75^. For each frequency band, bandpass filtering was performed by sequentially applying a high-pass and a low-pass FIR filter, with passband edges defined at the -6 dB cutoff frequencies of the respective filters. To avoid phase delays from the filtering process, we applied zero-phase filters. Following bandpass filtering, the instantaneous phase of the LFP at each time point was computed as the angle of the analytic signal obtained via the Hilbert transform within each frequency band.

### Direction tuning measurement and estimation from LFP signals

Before the smooth pursuit eye movement task, the direction tuning of each channel recorded in area MT was estimated in a separate direction tuning task. To assess the direction tuning of the recording channel, the stimulus motion was presented in 12 directions (0 to 330° in 30° increments) at the preferred speed of the recording channels while the animals fixated their eye gaze on a central stationary spot. A circular patch of random dots with 100% coherence served as the visual stimulus. Each trial began with a fixation duration between 600 and 1200 ms, and then six different motion stimuli were presented for 256 ms with 300 ms of interposed stationary epochs. Here, we approximate the overall direction tuning of local neural population through calculating LFP amplitude at high-gamma frequency band (70-150 Hz) following a previous study suggesting tight relationship between high-gamma LFP and multiunit activity of local neural population^36-38^. We bandpass-filtered the LFP at the high-gamma frequency band and applied Hilber transformation to obtain the amplitude. The LFP amplitude was then averaged over the 50-306 ms following stimulus onset. The preferred direction of the given LFP channel was determined by the direction tuning responses in high contrast condition, where the stimulus direction (among the 12 directions) that induced the maximum response amplitude was selected as the preferred direction. The estimated preferred directions obtained from this process were used in the analyses aimed at observing the power and phase changes as a function of the difference between the preferred direction of each channel and the prior direction.

### Statistical test

To address the multiple comparisons problem in group comparisons for LFP power across frequencies, we employed a two-tailed, paired two-sample *t*-test combined with a nonparametric, cluster-based permutation test, implemented using a statistical toolbox in MATLAB^77^. Since comparisons were made between two conditions (*prior vs. no prior*) within the same set of recording channels, a paired statistical test was appropriate. Assuming that neighboring frequency points are correlated, significant frequency clusters were defined as contiguous frequency points where the uncorrected *p*-values from *t*-test were < 0.05. To assess the statistical significance of each frequency cluster, we generated a null distribution of cluster-level statistics by randomly shuffling the condition labels across trials (10,000 permutations). For each permutation, *t*-values were computed, and the maximum cluster-level sum of *t*-values was extracted to build the null distribution. Observed clusters were considered statistically significant if their summed *t*-values exceeded the 95th percentile of the null distribution (cluster-level corrected *p* < 0.05). To investigate whether the LFP power (or phase) reduction due to prior expectations was associated with Δθ (the absolute difference between the prior direction and each neuron’s preferred direction), we performed linear regression analysis. Regression coefficients were computed between Δθ and the LFP power difference (or phase difference) between the *prior* and *no prior* conditions.

## Acknowledgement

This research was supported by the National Research Foundation of Korea grant funded by the Korean government (MSIT) (RS-2023-NR077084, RS-2023-00217361, RS-2025-00518788, and RS-2025-02263832).

## References

1. Fetsch, C.R., Pouget, A., DeAngelis, G.C., and Angelaki, D.E. (2011). Neural correlates of reliability-based cue weighting during multisensory integration. Nat Neurosci 15, 146–154. 10.1038/nn.2983.

2. Funamizu, A., Kuhn, B., and Doya, K. (2016). Neural substrate of dynamic Bayesian inference in the cerebral cortex. Nat Neurosci 19, 1682–1689. 10.1038/nn.4390.

3. Jeong, W., Kim, S., Park, J., and Lee, J. (2023). Multivariate EEG activity reflects the Bayesian integration and the integrated Galilean relative velocity of sensory motion during sensorimotor behavior. Commun Biol 6, 113. 10.1038/s42003-023-04481-2.

4. Kording, K.P., and Wolpert, D.M. (2004). Bayesian integration in sensorimotor learning. Nature 427, 244–247. 10.1038/nature02169.

5. Zhang, L.Q., and Stocker, A.A. (2022). Prior Expectations in Visual Speed Perception Predict Encoding Characteristics of Neurons in Area MT. J Neurosci 42, 2951–2962. 10.1523/JNEUROSCI.1920-21.2022.

6. Akrami, A., Kopec, C.D., Diamond, M.E., and Brody, C.D. (2018). Posterior parietal cortex represents sensory history and mediates its effects on behaviour. Nature 554, 368–372. 10.1038/nature25510.

7. Findling, C., Hubert, F., International Brain, L., Acerbi, L., Benson, B., Benson, J., Birman, D., Bonacchi, N., Buchanan, E.K., Bruijns, S., et al. (2025). Brain-wide representations of prior information in mouse decision-making. Nature 645, 192–200. 10.1038/s41586-025-09226-1.

8. Ishizu, K., Nishimoto, S., Ueoka, Y., and Funamizu, A. (2024). Localized and global representation of prior value, sensory evidence, and choice in male mouse cerebral cortex. Nat Commun 15, 4071. 10.1038/s41467-024-48338-6.

9. Langlois, T.A., Charlton, J.A., and Goris, R.L.T. (2025). Bayesian inference by visuomotor neurons in the prefrontal cortex. Proc Natl Acad Sci U S A 122, e2420815122. 10.1073/pnas.2420815122.

10. Sohn, H., Narain, D., Meirhaeghe, N., and Jazayeri, M. (2019). Bayesian Computation through Cortical Latent Dynamics. Neuron 103, 934–947 e935. 10.1016/j.neuron.2019.06.012.

11. Gonzalez-Garcia, C., and He, B.J. (2021). A Gradient of Sharpening Effects by Perceptual Prior across the Human Cortical Hierarchy. J Neurosci 41, 167–178. 10.1523/JNEUROSCI.2023-20.2020.

12. Ogando, M.B., Abdeladim, L., Sit, K.K., Shin, H., Sridharan, S., Gopakumar, K., and Adesnik, H. (2025). Feature-specific inhibitory connectivity augments the accuracy of cortical representations. bioRxiv. 10.1101/2025.08.02.668307.

13. Park, J., Kim, S., Kim, H.R., and Lee, J. (2023). Prior expectation enhances sensorimotor behavior by modulating population tuning and subspace activity in sensory cortex. Sci Adv 9, eadg4156. 10.1126/sciadv.adg4156.

14. Driscoll, L.N., Shenoy, K., and Sussillo, D. (2024). Flexible multitask computation in recurrent networks utilizes shared dynamical motifs. Nat Neurosci 27, 1349–1363. 10.1038/s41593-024-01668-6.

15. Song, H.F., Yang, G.R., and Wang, X.J. (2016). Training Excitatory-Inhibitory Recurrent Neural Networks for Cognitive Tasks: A Simple and Flexible Framework. PLoS Comput Biol 12, e1004792. 10.1371/journal.pcbi.1004792.

16. Yang, G.R., and Molano-Mazon, M. (2021). Towards the next generation of recurrent network models for cognitive neuroscience. Curr Opin Neurobiol 70, 182–192. 10.1016/j.conb.2021.10.015.

17. Darlington, T.R., Beck, J.M., and Lisberger, S.G. (2018). Neural implementation of Bayesian inference in a sensorimotor behavior. Nat Neurosci 21, 1442–1451. 10.1038/s41593-018-0233-y.

18. Yang, J., Lee, J., and Lisberger, S.G. (2012). The interaction of bayesian priors and sensory data and its neural circuit implementation in visually guided movement. J Neurosci 32, 17632–17645. 10.1523/JNEUROSCI.1163-12.2012.

19. Michaels, J.A., Dann, B., and Scherberger, H. (2016). Neural Population Dynamics during Reaching Are Better Explained by a Dynamical System than Representational Tuning. PLoS Comput Biol 12, e1005175. 10.1371/journal.pcbi.1005175.

20. Raiguel, S.E., Xiao, D.K., Marcar, V.L., and Orban, G.A. (1999). Response latency of macaque area MT/V5 neurons and its relationship to stimulus parameters. J Neurophysiol 82, 1944–1956. 10.1152/jn.1999.82.4.1944.

21. Stroud, J.P., Porter, M.A., Hennequin, G., and Vogels, T.P. (2018). Motor primitives in space and time via targeted gain modulation in cortical networks. Nat Neurosci 21, 1774–1783. 10.1038/s41593-018-0276-0.

22. Chelaru, M.I., and Dragoi, V. (2008). Efficient coding in heterogeneous neuronal populations. Proc Natl Acad Sci U S A 105, 16344–16349. 10.1073/pnas.0807744105.

23. Gast, R., Solla, S.A., and Kennedy, A. (2024). Neural heterogeneity controls computations in spiking neural networks. Proc Natl Acad Sci U S A 121, e2311885121. 10.1073/pnas.2311885121.

24. Mejias, J.F., and Longtin, A. (2012). Optimal heterogeneity for coding in spiking neural networks. Phys Rev Lett 108, 228102. 10.1103/PhysRevLett.108.228102.

25. Perez-Nieves, N., Leung, V.C.H., Dragotti, P.L., and Goodman, D.F.M. (2021). Neural heterogeneity promotes robust learning. Nat Commun 12, 5791. 10.1038/s41467-021-26022-3.

26. Mante, V., Sussillo, D., Shenoy, K.V., and Newsome, W.T. (2013). Context-dependent computation by recurrent dynamics in prefrontal cortex. Nature 503, 78–84. 10.1038/nature12742.

27. Kozyrev, V., Daliri, M.R., Schwedhelm, P., and Treue, S. (2019). Strategic deployment of feature-based attentional gain in primate visual cortex. PLoS Biol 17, e3000387. 10.1371/journal.pbio.3000387.

28. Treue, S., and Martinez Trujillo, J.C. (1999). Feature-based attention influences motion processing gain in macaque visual cortex. Nature 399, 575–579. 10.1038/21176.

29. Womelsdorf, T., Anton-Erxleben, K., and Treue, S. (2008). Receptive field shift and shrinkage in macaque middle temporal area through attentional gain modulation. J Neurosci 28, 8934–8944. 10.1523/JNEUROSCI.4030-07.2008.

30. Wimmer, K., Nykamp, D.Q., Constantinidis, C., and Compte, A. (2014). Bump attractor dynamics in prefrontal cortex explains behavioral precision in spatial working memory. Nat Neurosci 17, 431–439. 10.1038/nn.3645.

31. Compte, A., Brunel, N., Goldman-Rakic, P.S., and Wang, X.J. (2000). Synaptic mechanisms and network dynamics underlying spatial working memory in a cortical network model. Cereb Cortex 10, 910–923. 10.1093/cercor/10.9.910.

32. Sanchez-Vives, M.V., and McCormick, D.A. (2000). Cellular and network mechanisms of rhythmic recurrent activity in neocortex. Nat Neurosci 3, 1027–1034. 10.1038/79848.

33. Saleem, A.B., Chadderton, P., Apergis-Schoute, J., Harris, K.D., and Schultz, S.R. (2010). Methods for predicting cortical UP and DOWN states from the phase of deep layer local field potentials. J Comput Neurosci 29, 49–62. 10.1007/s10827-010-0228-5.

34. Okun, M., Naim, A., and Lampl, I. (2010). The subthreshold relation between cortical local field potential and neuronal firing unveiled by intracellular recordings in awake rats. J Neurosci 30, 4440–4448. 10.1523/JNEUROSCI.5062-09.2010.

35. Tan, A.Y., Chen, Y., Scholl, B., Seidemann, E., and Priebe, N.J. (2014). Sensory stimulation shifts visual cortex from synchronous to asynchronous states. Nature 509, 226–229. 10.1038/nature13159.

36. Liu, J., and Newsome, W.T. (2006). Local field potential in cortical area MT: stimulus tuning and behavioral correlations. J Neurosci 26, 7779–7790. 10.1523/JNEUROSCI.5052-05.2006.

37. Ray, S., Crone, N.E., Niebur, E., Franaszczuk, P.J., and Hsiao, S.S. (2008). Neural correlates of high-gamma oscillations (60-200 Hz) in macaque local field potentials and their potential implications in electrocorticography. J Neurosci 28, 11526–11536. 10.1523/JNEUROSCI.2848-08.2008.

38. Ray, S., and Maunsell, J.H. (2011). Different origins of gamma rhythm and high-gamma activity in macaque visual cortex. PLoS Biol 9, e1000610. 10.1371/journal.pbio.1000610.

39. Kok, P., Brouwer, G.J., van Gerven, M.A., and de Lange, F.P. (2013). Prior expectations bias sensory representations in visual cortex. J Neurosci 33, 16275–16284. 10.1523/JNEUROSCI.0742-13.2013.

40. Foeller, E., Celikel, T., and Feldman, D.E. (2005). Inhibitory sharpening of receptive fields contributes to whisker map plasticity in rat somatosensory cortex. J Neurophysiol 94, 4387–4400. 10.1152/jn.00553.2005.

41. Lee, S.H., Kwan, A.C., Zhang, S., Phoumthipphavong, V., Flannery, J.G., Masmanidis, S.C., Taniguchi, H., Huang, Z.J., Zhang, F., Boyden, E.S., et al. (2012). Activation of specific interneurons improves V1 feature selectivity and visual perception. Nature 488, 379–383. 10.1038/nature11312.

42. Li, Y.T., Ma, W.P., Li, L.Y., Ibrahim, L.A., Wang, S.Z., and Tao, H.W. (2012). Broadening of inhibitory tuning underlies contrast-dependent sharpening of orientation selectivity in mouse visual cortex. J Neurosci 32, 16466–16477. 10.1523/JNEUROSCI.3221-12.2012.

43. Ghose, G.M., and Maunsell, J.H. (2008). Spatial summation can explain the attentional modulation of neuronal responses to multiple stimuli in area V4. J Neurosci 28, 5115–5126. 10.1523/JNEUROSCI.0138-08.2008.

44. Martinez-Trujillo, J., and Treue, S. (2002). Attentional modulation strength in cortical area MT depends on stimulus contrast. Neuron 35, 365–370. 10.1016/s0896-6273(02)00778-x.

45. Maunsell, J.H., and Treue, S. (2006). Feature-based attention in visual cortex. Trends Neurosci 29, 317–322. 10.1016/j.tins.2006.04.001.

46. Watson, J.F., Ho, H., and Greger, I.H. (2017). Synaptic transmission and plasticity require AMPA receptor anchoring via its N-terminal domain. Elife 6. 10.7554/eLife.23024.

47. Gainey, M.A., Hurvitz-Wolff, J.R., Lambo, M.E., and Turrigiano, G.G. (2009). Synaptic scaling requires the GluR2 subunit of the AMPA receptor. J Neurosci 29, 6479–6489. 10.1523/JNEUROSCI.3753-08.2009.

48. Blanke, M.L., and VanDongen, A.M.J. (2009). Activation Mechanisms of the NMDA Receptor. In Biology of the NMDA Receptor, A.M. Van Dongen, ed.

49. Huang, X., Sun, X., Wang, Q., Zhang, J., Wen, H., Chen, W.J., and Zhu, S. (2025). Structural insights into the diverse actions of magnesium on NMDA receptors. Neuron 113, 1006–1018 e1004. 10.1016/j.neuron.2025.01.021.

50. Sherwood, M.W., Oliet, S.H.R., and Panatier, A. (2021). NMDARs, Coincidence Detectors of Astrocytic and Neuronal Activities. Int J Mol Sci 22. 10.3390/ijms22147258.

51. Bessieres, B., Dupuis, J., Groc, L., Bontempi, B., and Nicole, O. (2024). Synaptic rearrangement of NMDA receptors controls memory engram formation and malleability in the cortex. Sci Adv 10, eado1148. 10.1126/sciadv.ado1148.

52. Hunt, D.L., and Castillo, P.E. (2012). Synaptic plasticity of NMDA receptors: mechanisms and functional implications. Curr Opin Neurobiol 22, 496–508. 10.1016/j.conb.2012.01.007.

53. Jain, V., Murphy-Baum, B.L., deRosenroll, G., Sethuramanujam, S., Delsey, M., Delaney, K.R., and Awatramani, G.B. (2020). The functional organization of excitation and inhibition in the dendrites of mouse direction-selective ganglion cells. Elife 9. 10.7554/eLife.52949.

54. Nadim, F., and Bucher, D. (2014). Neuromodulation of neurons and synapses. Curr Opin Neurobiol 29, 48–56. 10.1016/j.conb.2014.05.003.

55. Wilson, D.E., Scholl, B., and Fitzpatrick, D. (2018). Differential tuning of excitation and inhibition shapes direction selectivity in ferret visual cortex. Nature 560, 97–101. 10.1038/s41586-018-0354-1.

56. Arnal, L.H., and Giraud, A.L. (2012). Cortical oscillations and sensory predictions. Trends Cogn Sci 16, 390–398. 10.1016/j.tics.2012.05.003.

57. Son, S., Moon, J., Kim, Y.J., Kang, M.S., and Lee, J. (2023). Frontal-to-visual information flow explains predictive motion tracking. Neuroimage 269, 119914. 10.1016/j.neuroimage.2023.119914.

58. Bastos, A.M., Vezoli, J., Bosman, C.A., Schoffelen, J.M., Oostenveld, R., Dowdall, J.R., De Weerd, P., Kennedy, H., and Fries, P. (2015). Visual areas exert feedforward and feedback influences through distinct frequency channels. Neuron 85, 390–401. 10.1016/j.neuron.2014.12.018.

59. Buschman, T.J., and Miller, E.K. (2007). Top-down versus bottom-up control of attention in the prefrontal and posterior parietal cortices. Science 315, 1860–1862. 10.1126/science.1138071.

60. Siegel, M., Donner, T.H., Oostenveld, R., Fries, P., and Engel, A.K. (2007). High-frequency activity in human visual cortex is modulated by visual motion strength. Cereb Cortex 17, 732–741. 10.1093/cercor/bhk025.

61. Aliramezani, M., Singh, B., Constantinidis, C., and Daliri, M.R. (2025). Low-frequency local field potentials reveal integration of spatial and non-spatial information in prefrontal cortex. Neuroimage 310, 121172. 10.1016/j.neuroimage.2025.121172.

62. Bastos, G., Holmes, J.T., Ross, J.M., Rader, A.M., Gallimore, C.G., Wargo, J.A., Peterka, D.S., and Hamm, J.P. (2023). Top-down input modulates visual context processing through an interneuron-specific circuit. Cell Rep 42, 113133. 10.1016/j.celrep.2023.113133.

63. Samaha, J., Bauer, P., Cimaroli, S., and Postle, B.R. (2015). Top-down control of the phase of alpha-band oscillations as a mechanism for temporal prediction. Proc Natl Acad Sci U S A 112, 8439–8444. 10.1073/pnas.1503686112.

64. Clark, D.G., and Beiran, M. (2025). Structure of activity in multiregion recurrent neural networks. Proc Natl Acad Sci U S A 122, e2404039122. 10.1073/pnas.2404039122.

65. Shenoy, K.V., and Kao, J.C. (2021). Measurement, manipulation and modeling of brain-wide neural population dynamics. Nat Commun 12, 633. 10.1038/s41467-020-20371-1.

66. Kang, B., and Druckmann, S. (2020). Approaches to inferring multi-regional interactions from simultaneous population recordings: Inferring multi-regional interactions from simultaneous population recordings. Curr Opin Neurobiol 65, 108–119. 10.1016/j.conb.2020.10.004.

67. Machado, T.A., Kauvar, I.V., and Deisseroth, K. (2022). Multiregion neuronal activity: the forest and the trees. Nat Rev Neurosci 23, 683–704. 10.1038/s41583-022-00634-0.

68. Lisberger, S.G., Morris, E.J., and Tychsen, L. (1987). Visual motion processing and sensory-motor integration for smooth pursuit eye movements. Annu Rev Neurosci 10, 97–129. 10.1146/annurev.ne.10.030187.000525.

69. Lee, J., and Lisberger, S.G. (2013). Gamma synchrony predicts neuron-neuron correlations and correlations with motor behavior in extrastriate visual area MT. J Neurosci 33, 19677–19688. 10.1523/JNEUROSCI.3478-13.2013.

70. Gertz, E.M., Hiekkalinna, T., Digabel, S.L., Audet, C., Terwilliger, J.D., and Schaffer, A.A. (2014). PSEUDOMARKER 2.0: efficient computation of likelihoods using NOMAD. BMC Bioinformatics 15, 47. 10.1186/1471-2105-15-47.

71. Sussillo, D., and Abbott, L.F. (2009). Generating coherent patterns of activity from chaotic neural networks. Neuron 63, 544–557. 10.1016/j.neuron.2009.07.018.

72. Kim, R., Li, Y., and Sejnowski, T.J. (2019). Simple framework for constructing functional spiking recurrent neural networks. Proc Natl Acad Sci U S A 116, 22811–22820. 10.1073/pnas.1905926116.

73. Schwartz, A.B., Kettner, R.E., and Georgopoulos, A.P. (1988). Primate motor cortex and free arm movements to visual targets in three-dimensional space. I. Relations between single cell discharge and direction of movement. J Neurosci 8, 2913–2927. 10.1523/JNEUROSCI.08-08-02913.1988.

74. Georgopoulos, A.P., Kettner, R.E., and Schwartz, A.B. (1988). Primate motor cortex and free arm movements to visual targets in three-dimensional space. II. Coding of the direction of movement by a neuronal population. J Neurosci 8, 2928–2937. 10.1523/JNEUROSCI.08-08-02928.1988.

75. Delorme, A., and Makeig, S. (2004). EEGLAB: an open source toolbox for analysis of single-trial EEG dynamics including independent component analysis. J Neurosci Methods 134, 9–21. 10.1016/j.jneumeth.2003.10.009.

76. Bokil, H., Andrews, P., Kulkarni, J.E., Mehta, S., and Mitra, P.P. (2010). Chronux: a platform for analyzing neural signals. J Neurosci Methods 192, 146–151. 10.1016/j.jneumeth.2010.06.020.

77. Maris, E., and Oostenveld, R. (2007). Nonparametric statistical testing of EEG- and MEG-data. J Neurosci Methods 164, 177–190. 10.1016/j.jneumeth.2007.03.024.

